# The developmental timing of spinal touch processing alterations and its relation to ASD-associated behaviors in mouse models

**DOI:** 10.1101/2023.05.09.539589

**Authors:** Aniqa Tasnim, Ilayda Alkislar, Richard Hakim, Josef Turecek, Amira Abdelaziz, Lauren L. Orefice, David D. Ginty

## Abstract

Altered somatosensory reactivity is frequently observed among individuals with autism spectrum disorders (ASDs). Here, we report that while multiple mouse models of ASD exhibit aberrant somatosensory behaviors in adulthood, some models exhibit altered tactile reactivity as early as embryonic development, while in others, altered reactivity emerges later in life. Additionally, tactile over-reactivity during neonatal development is associated with anxiety-like behaviors and social interaction deficits in adulthood, whereas tactile over-reactivity that emerges later in life is not. The locus of circuit disruption dictates the timing of aberrant tactile behaviors: altered feedback or presynaptic inhibition of peripheral mechanosensory neurons leads to abnormal tactile reactivity during neonatal development, while disruptions in feedforward inhibition in the spinal cord lead to touch reactivity alterations that manifest later in life. Thus, the developmental timing of aberrant touch processing can predict the manifestation of ASD-associated behaviors in mouse models, and differential timing of sensory disturbance onset may contribute to phenotypic diversity across individuals with ASD.

## MAIN TEXT

Sensory processing dysfunction is reported in over 94% of children, adolescents, and adults with autism spectrum disorders (ASD)^1–4^, and the latest Diagnostic and Statistical Manual of Mental Disorders (DSM-V) includes atypical responses to sensory stimuli as a core diagnostic criterion for ASD^5^. Notably, 60% of individuals with ASD report tactile sensitivity difficulties^4^, with the initial appearance of atypical touch reactivity often predating ASD diagnoses^6–9^. Interacting with the world through the sense of touch, particularly at young ages, plays a central role in cognitive and social development in humans, non-human primates, and rodents^10–13^, and early alterations to tactile processing and reactivity can be predictive of the emergence and severity of ASD-associated traits later in life^6, 14, 15^. However, autism diagnoses are elusive during infancy, in part due to high variability in ASD phenotypic presentation across individuals^16^. Defining atypical sensory processing behaviors during early development as they relate to other core features of ASD may aid in early detection, diagnosis, and prognosis.

The heterogeneity in ASD etiology and behavioral outcomes necessitates a greater focus on defining neurophysiological signatures that may define subgroups in ASD^17^. Moreover, understanding neurophysiological changes in touch processing in ASD will inform the design of therapeutic approaches to treat sensory symptoms, which may in turn mitigate other ASD-related symptoms in certain individuals^12^. Recent studies in genetic models of ASD have revealed that rodents harboring mutations in the ASD-associated genes *Mecp2*, *Gabrb3*, *Shank3*, and *Fmr1* exhibit over-reactivity to touch stimuli^12, 18–24^. Peripheral mechanosensory neurons, which convey information to the spinal cord and brainstem about tactile stimuli acting on the skin, cell-autonomously require proper expression of *Mecp2*, *Gabrb3*, and *Shank3* for normal tactile reactivity in adulthood^12, 23, 25^. Remarkably, developmental deletion of these genes only from peripheral somatosensory neurons leads to abnormal tactile behaviors, deficits in social interactions, and increased anxiety-like behaviors in adulthood. However, genetic ablation of these genes late in postnatal development drives tactile over-reactivity, but does not lead to changes in social or anxiety-like behaviors^12^. Thus, in at least some models of ASD, somatosensory deficits that arise in the periphery during development contribute to the generation of certain ASD-associated behaviors.

Here, we sought to understand how the loci of molecular and circuit disruption in disparate ASD mouse models may predict altered touch reactivity and other ASD-related behavioral manifestations in adulthood. We found that tactile behavioral alterations are observed across mutant models in adulthood, but that disruptions to tactile reactivity emerge at different developmental times. Genetic models with dysfunction in presynaptic, feedback inhibition of peripheral sensory neurons exhibit touch over-reactivity perinatally, and display anxiety-like and social interaction deficits in adulthood. Conversely, genetic models with disruption to axo-dendritic/somatic feedforward inhibition in spinal touch circuits have normal touch reactivity neonatally and enhanced tactile reactivity later in life. However, this elevation of tactile reactivity that manifests later in life occurs in the absence of anxiety-like and social behavioral deficits. Our findings highlight neurophysiological mechanisms that underlie susceptibility to aberrant touch processing in ASD models, and we show that early over-reactivity is predictive of abnormal anxiety-like and social behaviors in adulthood. Thus, differences in the developmental timing of aberrant sensory reactivity onset may contribute to the heterogeneity of core and co-morbid ASD-associated behaviors.

## RESULTS

### Multiple genetic ASD mouse models exhibit tactile over-reactivity, with differences in cognitive and social ASD-related behaviors

ASD-associated genes *Mecp2* and *Gabrb3* are required in peripheral somatosensory neurons for proper axo-axonic (presynaptic) feedback inhibition of these neurons and normal tactile behaviors in mice^22, 23^. Consistent with previous findings, adult animals heterozygous for *Gabrb3* (*Gabrb3*^+/-^) or hemizygous for *Mecp2* (*Mecp2*^-/y^) exhibited enhanced sensitivity to back hairy skin stimulation, measured by tactile prepulse inhibition of an acoustic startle response (tactile PPI) and to a gentle air puff stimulus alone (Fig. 1a-c)^23^. Moreover, these animals exhibited increased anxiety-like behaviors, measured by a reduction of time spent in the center of a chamber during the open field test (Fig. 1d,e), as well as abnormal social interaction behaviors, measured by the duration of interaction with a novel mouse compared to an empty cup during the three-chamber social interaction test (Fig. 1f,g)^23, 24, 26–28^. To explore conserved and additional mechanisms of altered sensory reactivity in mouse models of ASD, we asked whether other ASD-associated genes implicated in inhibitory synaptic signaling are also required for tactile behaviors.

**Figure 1.**
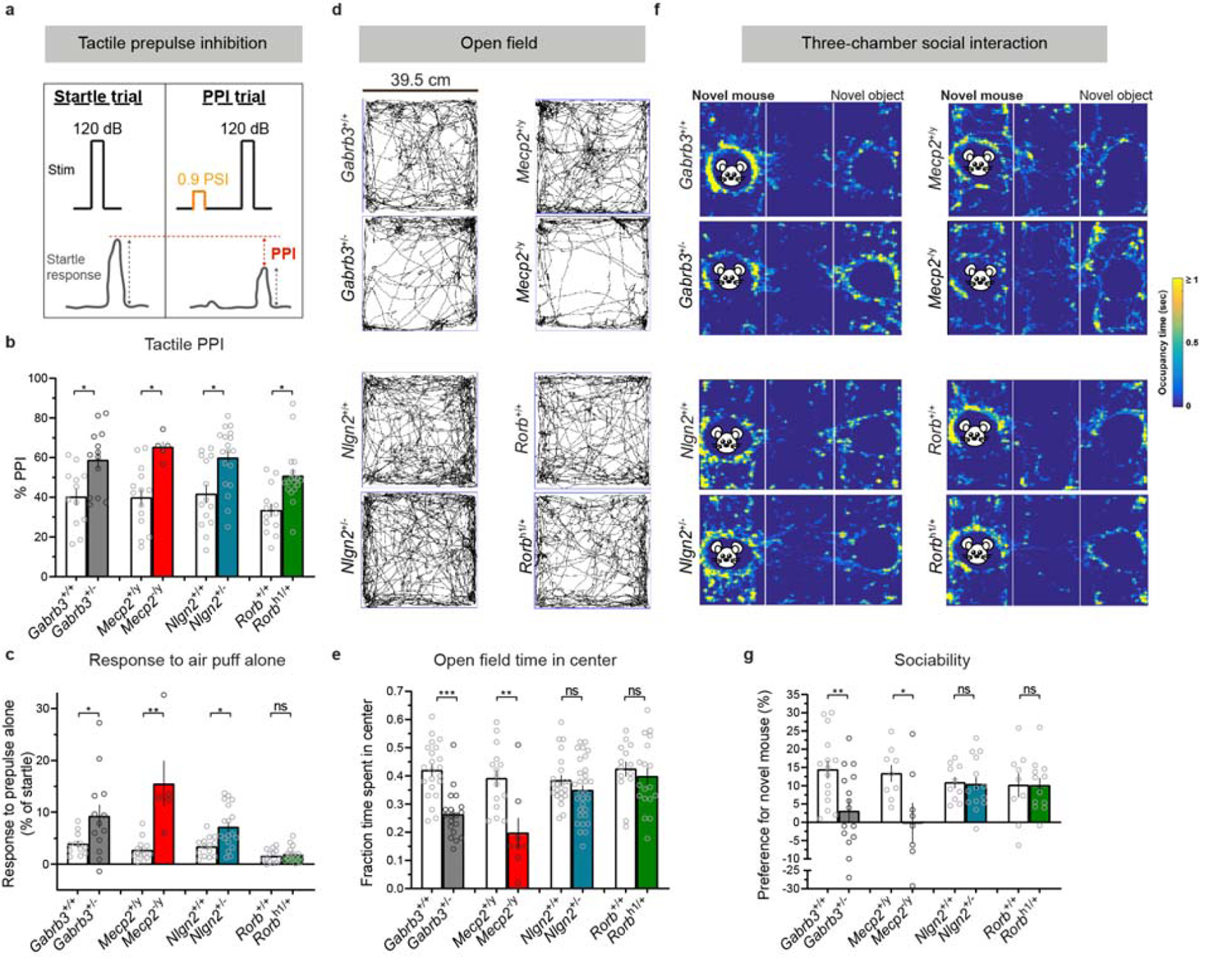
Multiple global knockout models of ASD-associated genes exhibit tactile over-reactivity in adulthood, with differences in ASD-associated behaviors. **a**, Diagram showing tactile prepulse inhibition of the startle reflex assay paradigm. The decrease in the startle response to a 120 dB acoustic pulse when preceded by a light air puff is defined as the effect of PPI (measured as a percent). The interval between the prepulse and pulse is 250 milliseconds. **b**, Tactile PPI. *P*=0.0071 for *Gabrb3*, *P*=0.0033 for *Mecp2*, *P*=0.0043 for *Nlgn2*, *P*=0.0015 for *Rorb*, unpaired two-tailed t-tests. **c**, Response to the 0.9 PSI air puff stimulus alone, expressed as a percent of startle response to an acoustic startle stimulus. *P*=0.0457 for *Gabrb3*, *P*=0.0003 for *Mecp2, P*=0.0054 for *Nlgn2*, *P*=0.7374 for *Rorb*, Mann-Whitney *U* tests. **d**, Representative activity traces in the open field assay for mutant mice and control littermates. **e**, Fraction of time spent in the center of the open field chamber. *P*<0.0001 for *Gabrb3*, *P*=0.2050 for *Nlgn2*, *P*=0.5339 for *Rorb*, unpaired two-tailed t-tests. **f**, Representative heat maps showing time spent with a novel mouse or novel object (an empty cup) in the 3-chamber social interaction assay for mutant mice and control littermates. **g**, Percent preference for a novel mouse over a novel object in the 3-chamber social interaction assay. *P*=0.0091 for *Gabrb3*, *P*=0.8603 for *Nlgn2*, *P*=0.9887 for *Rorb*, unpaired two-tailed t-tests. For **b, c, e,** and **g,** data represent means±s.e.m.

The ASD-associated genes *Nlgn2* and *Nlgn4* encode the synaptic-adhesion molecules Neuroligin-2 and Neuroligin-4, respectively, which contribute to GABA_A_ and glycine receptor synapse function in the forebrain and brainstem^29–32^. We observed that animals harboring loss-of-function mutations in *Nlgn2*^33^ exhibited increased responsivity to hairy skin stimulation (Fig. 1b,c). Additionally, *Nlgn2*^+/-^ animals displayed deficits in a texture discrimination assay, but exhibited normal preferences for novel objects differing in shape and color, indicating intact novelty preference (Extended Data Fig. 1a,b). Despite these marked changes in tactile reactivity and texture discrimination, *Nlgn2*^+/-^ mice showed no alterations in anxiety-like or social interaction behaviors compared to control littermates (Fig. 1d-g), while *Nlgn2*^-/-^ mice exhibited a partial increased anxiety-like phenotype, consistent with previous reports (Extended Data Fig. 1c,d)^34, 35^. In contrast, *Nlgn4* mutant animals^36^ did not exhibit detectable tactile or anxiety-like behavioral alterations (Extended Data Fig. 1e-g), and prior studies observed intact social behaviors in these mutant animals^37^, which may be consistent with significant evolutionary divergence of *Nlgn4* in the murine genome from other mammalian NLGN4 genes^38, 39^.

We also examined the ASD-associated gene, *Rorb*^40^, which is expressed by a population of spinal cord interneurons that mediate axo-dendritic/somatic feedforward inhibition of spinal projection neurons in the deep dorsal horn^41^. The activity of *Rorb*-expressing interneurons is required for normal tactile and sensorimotor function^41, 42^. To investigate the role of the *Rorb* gene in ASD-associated behaviors, we tested animals with a pathogenic mutation arising at the *Rorb* locus (*Rorb*^h1^)^43, 44^. As observed in *Gabrb3*^+/-^, *Mecp2*^-/y^, and *Nlgn2*^+/-^ animals, *Rorb*^h1/+^ mice exhibited enhanced tactile sensitivity in the tactile PPI assay, although *Rorb*^h1/+^ mice did not have enhanced reactivity to a gentle air puff alone (Fig. 1b,c). Like *Nlgn2*^+/-^ animals, *Rorb* mutant mice showed normal anxiety-like and social behaviors when compared to control littermates (Fig. 1d-g and Extended Data Fig. 1d). Together, these findings reveal considerable heterogeneity across mouse models of ASD in adulthood: while *Gabrb3*^+/-^, *Mecp2*^-/y^, *Nlgn2*^+/-^, and *Rorb*^h1/+^ animals all exhibit aberrant tactile behaviors, *Gabrb3*^+/-^ and *Mecp2*^-/y^ animals also display enhanced anxiety-like behaviors and reduced sociability^23^, whereas *Nlgn2*^+/-^, and *Rorb*^h1/+^ animals do not.

### Tactile over-reactivity emerges at different developmental time points across different ASD models

Our results demonstrate that while tactile over-reactivity is a characteristic shared by multiple ASD mouse models in adulthood, there are differences in co-morbid ASD-like behaviors. Our prior work in *Gabrb3*, *Mecp2*, and *Shank3* ASD mouse models revealed a developmental window before postnatal day 10, during which normal tactile sensitivity is required for the formation of normal brain microcircuit properties and certain cognitive and social behaviors^12, 23^. Expression of these genes in somatosensory neurons of the dorsal root ganglia (DRG) was required during early postnatal development for normal tactile, anxiety-like, and social behaviors in adulthood. Thus, we hypothesized that tactile over-reactivity during neonatal and early postnatal life may deterime the extent to which mice display altered anxiety-like and social behaviors later in adulthood.

We asked when during development tactile over-reactivity emerges in the *Gabrb3*^+/-^, *Mecp2*^-/y^, *Nlgn2*^+/-^, and *Rorb*^h1/+^ mouse models. To address this, we assessed reactivity to light air puff stimuli applied to back hairy skin of early postnatal age mice. We chose a time point within the first postnatal week, postnatal day 4 (P4), to measure air puff reactivity. P4 was ideal for these measurements because we observed substantial hair shaft emergence across strains compared to younger pups, and relative quiescience on an experimental platform during interstimulus intervals compared to older pups. Air puffs delivered to back hairy skin of P4 pups resulted in animal displacement responses, and the amplitude of body displacement could be quantified using an optic flow analysis measurement (Fig. 2a,b and Supplementary Movie 1). In this assay, responsivity increased as the airpuff stimulus intensity increased, was dependent on hair deflection (Extended Data Fig. 2a), and was lost following topical application of lidocaine to back hairy skin to silence cutaneous sensory nerve fibers (Extended Data Fig. 2b).

**Figure 2.**
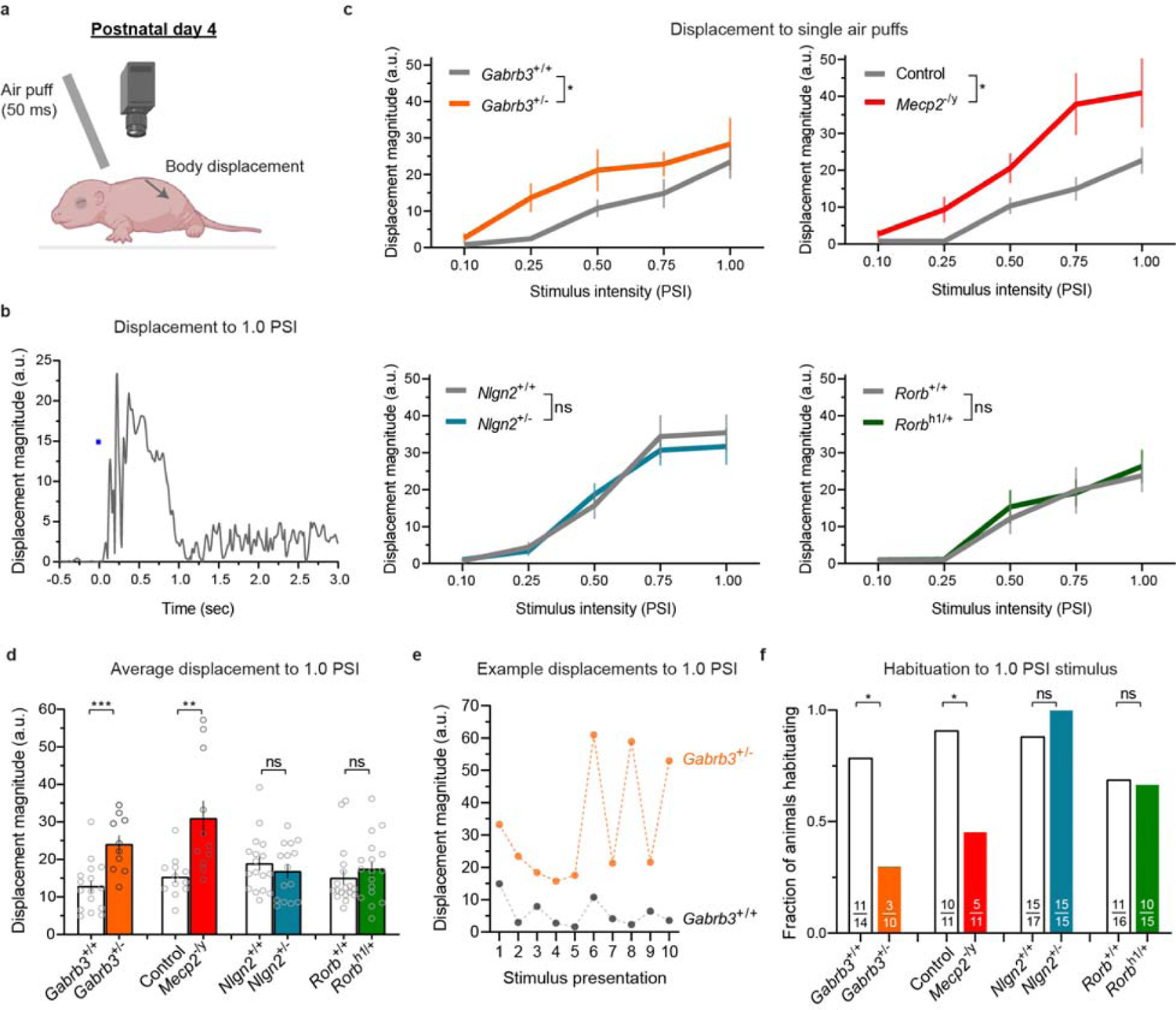
Neonatal tactile reactivity in global knockout ASD mouse models. **a**, Diagram showing experimental paradigm for neonatal air puff responsivity assay. Air puff tubing is affixed 3 mm above the nape of the neck of neonatal mice. Maximal body displacements in the 500 ms following a 50 ms air puff are measured using optic flow analysis. Parts created with <u>BioRender.com</u>. **b**, Example displacement trace of a control P4 mouse to a single 1.0 PSI air puff stimulus. **c**, Displacement responses to a single presentation of 0.10, 0.25, 0.50, 0.75 and 1.0 PSI stimuli. Two-way ANOVAs, effect of genotype; *Gabrb3*: (F[1, 24]=10.88, *P*=0.0030); *Mecp2*: (F[1, 21]=9.253, *P*=0.0062); *Nlgn2*: (F[1, 30]=0.1071, *P*=0.7457); *Rorb*: (F[1, 33]=0.1204, *P*=0.7308). Data represent means±s.e.m. **d**, Average displacement responses to 10 presentations of a 1.0 PSI stimulus at 20-30 second interstimulus intervals (ISIs). *P*=0.0003 for *Gabrb3*, *P*=0.0018 for *Mecp2*, *P*=0.2258 for *Nlgn2*, *P*=0.1970 for *Rorb*, unpaired one-tailed t-tests. Data represent means±s.e.m. **e**, Example traces showing displacements to ten presentations of a 1.0 PSI stimuli from a *Gabrb3*^+/+^ and *Gabrb3*^+/-^ mouse. To calculate habituation, the average of the last three stimulus presentation responses was compared to the average of the first three responses. A >25% reduction in responsivity between blocks was considered to be a habituation response. **f**, Fraction of animals habituating to repeated presentations of 1.0 PSI air puffs. For *Gabrb3, P*=0.0244; *Mecp2, P*=0.0317; *Nlgn2*, *P*=0.2742; *Rorb*, *P*=0.6017, one-sided Fisher’s Exact tests.

We hypothesized that *Gabrb3*^+/-^ and *Mecp2*^-/y^ animals, which exhibit enhanced anxiety-like behaviors and social interaction deficits in adulthood, would display aberrant tactile reactivity at this postnatal timepoint. In our experimental paradigm, we assessed reactivity to single presentations of 0.10, 0.25, 0.50, and 0.75 PSI air puff stimuli, followed by ten presentations of a 1.0 PSI stimulus, at 20-30 second intervals. Consistent with our hypothesis, P4 *Gabrb3*^+/-^ and *Mecp2*^-/y^ animals exhibited significantly enhanced reactivity to the individual presentations of the range of stimuli (Fig. 2c and Extended Data Fig. 2f) as well as to the repeated presentations 1.0 PSI stimulus (Fig. 2d). Additionally, we assessed habituation to the repeated 1.0 PSI air puff stimulus, as a lack of sensory habituation to repeated stimuli, including in the tactle domain, is frequently observed in individuals with or at high risk of ASD diagnoses^45–48^. To do so, we measured the decrease in response between the averages of the first and last three presentations of the 1.0 PSI stimulus (Fig. 2e), and categorized animals as habituating if they showed a >25% decrease in responsivity between these blocks. Interestingly, while the majority of control P4 pups exhibited habituation to repeated stimuli, a significantly smaller proportion of *Gabrb3*^+/-^ and *Mecp2*^-/y^ animals exhibited habituation (Fig. 2f). In immunohistochemical analyses, we confirmed that MeCP2 and GABRB3 proteins are indeed expressed perinatally, in both the DRG and the central axon terminals of DRG neurons, respectively (Extended Data Fig. 2c-e).

In contrast, and consistent with the hypothesis that normal tactile reactivity during development may be predictive of normal anxiety-like and social behaviors in adulthood, *Nlgn2* and *Rorb* mutant animals’ reactivity to air puff was indistinguishable from control littermates at P4 (Fig. 2c,d and Extended Data Fig. 2f,g). Moreover, these animals showed habituation that was comparable to control littermates (Fig. 2f and Extended Data Fig. 2h). Thus, aberrant tactile behaviors emerge at different times across the ASD mouse models tested here. Though *Nlgn2*^+/-^*, Rorb*^h1/+^, *Gabrb3*^+/-^, and *Mecp2*^-/y^ animals all display tactile over-reactivity in adulthood, only *Gabrb3*^+/-^ and *Mecp2*^-/y^ animals exhibit tactile over-reactivity at P4. *Nlgn2*^+/-^ and *Rorb*^h1/+^ animals are behaviorally indistinguishable from control littermates at this early postnatal time, but exhibit aberrant tactile behaviors in adulthood.

### Tactile over-reactivity is observed during embryonic development in *Gabrb3* mutant animals

Because a subset of ASD mutant models displayed tactile over-reactivity at P4, we next asked whether aberrant tactile behavior emerges even earlier in development. Additionally, we sought to test whether selective activation of mechanosensory neuron subtypes that mediate light touch would be sufficient to promote aberrant behavioral reactivity in mutant animals. To test this, we used optogenetic activation of low-threshold mechanoreceptor (LTMR) terminals in the skin and assessed the extent of behavioral reactivity to LTMR subtype-specific stimulation in postnatal day 0 (P0) mice (Fig. 3a).

**Figure 3.**
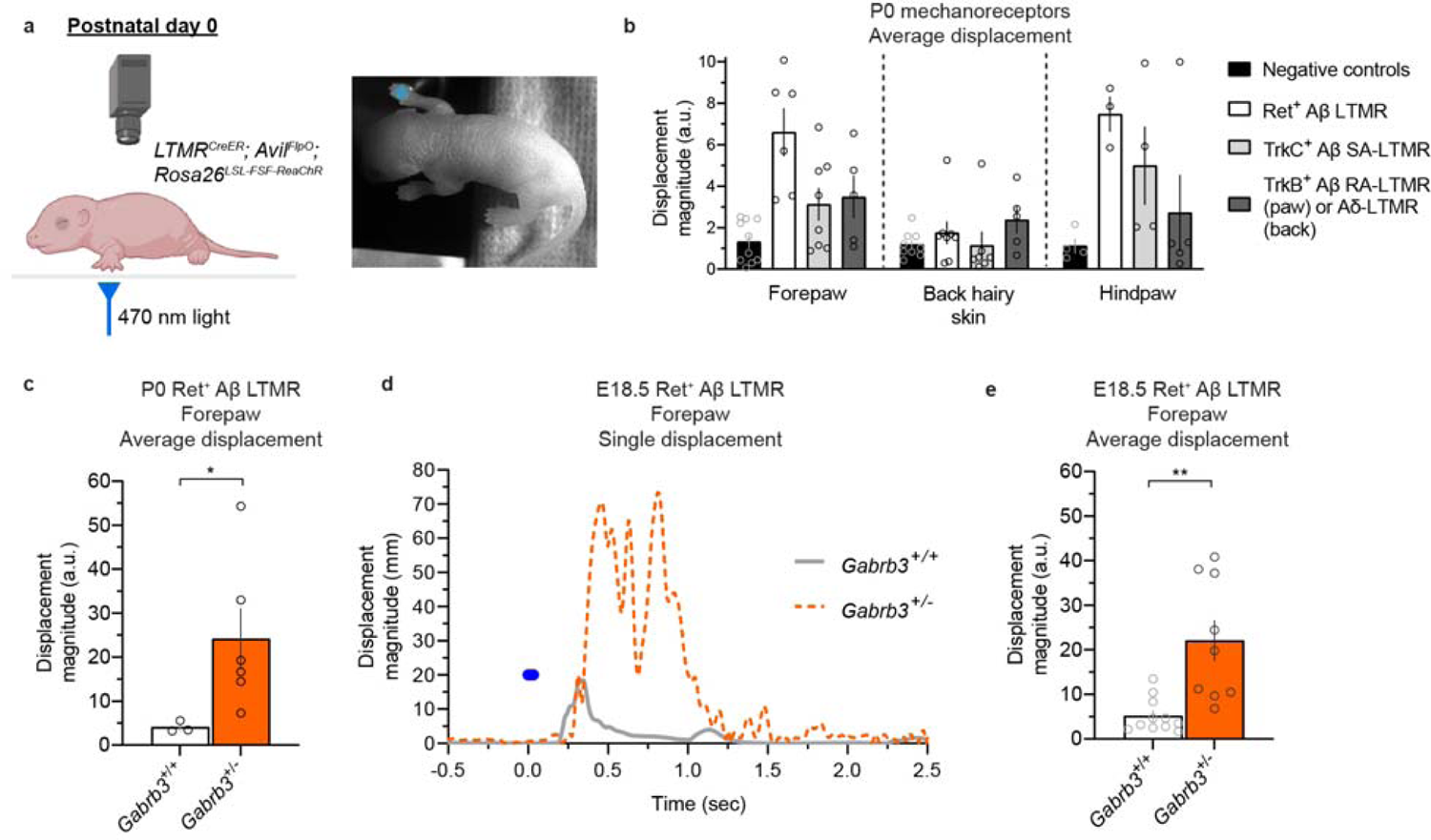
*Gabrb3* mutant mice are over-reactive to activation of A**β** low-threshold mechanoreceptors at birth and embryonic day 18.5. **a**, Experimental setup for perinatal mechanoreceptor optical activation assay. P0 or E18.5 mice are placed on a clear acrylic stage, and LED illumination is directed to the paw or back hairy skin. Parts created with <u>BioRender.com</u>. **b**, Average displacement responses to 5 optical stimuli (20-30 second ISIs) delivered to mechanoreceptor subtypes in forepaw, back hairy skin, and hindpaw compared to control littermates that lack opsin expression (controls include opsin-positive, Cre-negative and opsin-negative, Cre-positive animals). **c**, Average displacement responses to 5 optical stimuli activating Ret^+^ Aβ LTMRs to the forepaws of P0 *Ret^CreER^*; *Advillin^FlpO^*; *Rosa26^LSL-FSF-ReaChR-mCitrine^*; *Gabrb3*^+/+^ and *Gabrb3*^+/-^ animals. *P*=0.0169, unpaired one-tailed Welch’s t test. **d**, Example displacement traces from a E18.5 *Gabrb3*^+/+^ mouse and *Gabrb3*^+/-^ mouse to optical activation of Ret^+^ Aβ LTMRs in the forepaw. **e**, Average displacement responses to 5 optical stimuli activating Ret^+^ Aβ LTMRs in the forepaws of E18.5 control and mutant animals. *P*=0.0029, unpaired one-tailed Welch’s t test. For **b**, **c** and **e**, data represent means±s.e.m.

Using a Cre- and Flp-dependent ReaChR optogenetic actuator line, we first asked whether control P0 pups reacted to activation of large-diameter, LTMRs that terminate in the forepaw, hindpaw, and back hairy skin. Using the intersection of *Advillin^FlpO^* (to label all peripheral sensory neurons)^49^ and LTMR-specific CreER (LTMR*^CreER^*) lines, we targeted Ret^+^ Aβ LTMRs (using *Ret^CreER^*)^50^, Aβ slowly adapting LTMRs (Aβ SA-LTMRs, using *TrkC^CreER^*)^51^, and Aβ rapidly adapting LTMRs (Aβ RA-LTMRs in the glabrous paw, and Aδ-LTMRs in the hairy skin, using *TrkB^CreER^*)^52^ (Fig. 3b and Extended Data Fig. 3a). At this P0 time point, large-diameter sensory neurons, labeled by Neurofilament Heavy Chain (NFH), expressed ReaChR::mCitrine and innervated both glabrous and hairy skin (Extended Data Fig. 3a). We measured reactivity to stimulation using a 470 nm light pulse delivered five times at 20-30 second interstimulus intervals to the forepaw, back hairy skin, and hindpaw of P0 animals. We found that paw Ret^+^ Aβ LTMR stimulation evoked the greatest degree of reactivity (Fig. 3b). In contrast to behavioral responses evoked from the paw regions, activation of these same populations in back hairy skin resulted in less robust responses. We next tested forepaw activation of Ret^+^ Aβ LTMRs in *Gabrb3* mutants and their control littermates and found that mutant animals displayed enhanced responses to stimulation of Ret^+^ Aβ LTMRs at P0 (Fig. 3c). We also tested animals just prior to birth, on embryonic day 18.5 (E18.5). Reactivity could be reliably evoked from control E18.5 animals and, remarkably, the *Gabrb3* mutants were significantly more reactive to forepaw optical stimulation (Fig. 3d,e and Supplementary Movie 2), revealing a role for this ASD-associated gene in controlling tactile reactivity at late embryonic stages. Thus, selective activation of a subset of LTMRs innervating the paw is sufficient to evoke behavioral responses in E18.5 and P0 mice, and loss-of-function of *Gabrb3* can drive tactile behavioral over-reactivity at these ages.

### Differential cell-autonomous requirements for *Gabrb3* and *Nlgn2* in peripheral and spinal cord neuron types for tactile behaviors

Axo-axonic feedback inhibition, or presynaptic inhibition, is mediated by GABA_A_ receptors (GABA_A_Rs) present on peripheral somatosensory neuron afferent fibers, and deletion of *Gabrb3* or *Mecp2* in these neurons leads to a loss of GABA_A_R-mediated feedback inhibition^22, 23^. Consistent with a role in organizing both GABAergic and glycinergic synapses, the gene product of *Nlgn2* (Neuroligin-2, or NLGN2) was found to be required for both types of inhibitory neurotranmission in the medulla^30^. Interestingly, NLGN2 is found in native GABA_A_R complexes in the mouse brain^53^, and its transcripts are expressed in DRG neurons (Extended Data Fig. 4a). Therefore, we tested whether NLGN2 is involved in GABA_A_R scaffolding in sensory axon terminals. This was not the case, since GABA_A_R immunoreactivity associated with vesicular glutamate transporter 1 (VGLUT1)-positive sensory axon terminals in the dorsal horn of mice lacking NLGN2 in sensory neurons (*Advillin^Cre^*; *Nlgn2^flox/flox^* mice) was normal (Extended Data Fig. 4b,c), despite complete loss of *Nlgn2* expression in the DRGs of these mutants (Extended Data Fig. 4a). Thus, unlike GABRB3 and MeCP2, NLGN2 is not required in peripheral somatosensory neurons for GABA_A_R localization to their presynaptic terminals. Additionally, animals with either *Gabrb3* or *Mecp2* deleted from peripheral somatosensory neurons exhibited tactile over-reactivity in the tactile PPI assay and in response to an air puff alone, which was not the case in *Advillin^Cre^*; *Nlgn2^flox/flox^*mice (Fig. 4c,d and refs ^22, 23^), indicating that *Nlgn2* expression is not required in somatosensory neurons for tactile sensitivity.

**Figure 4.**
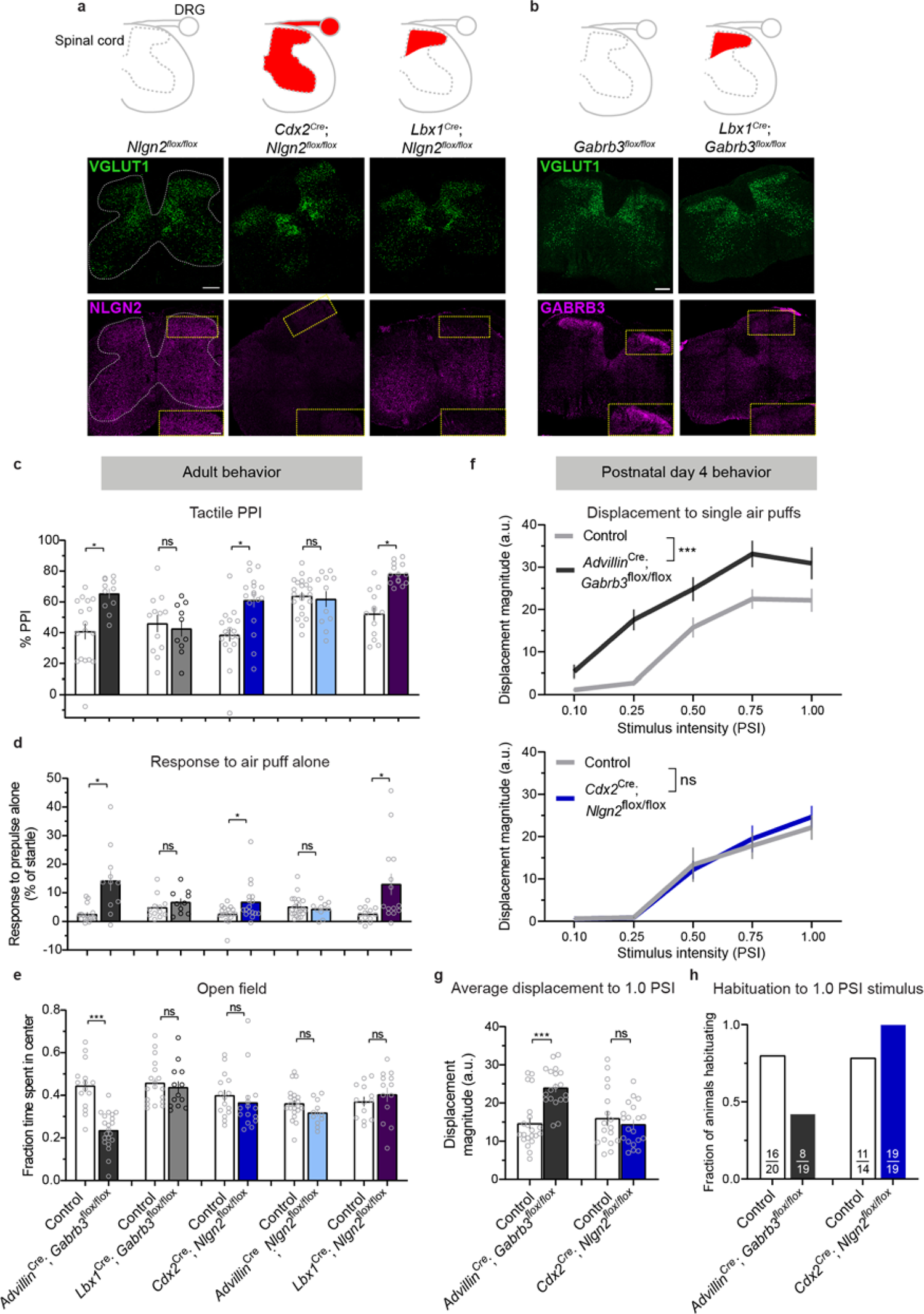
Differential cell-autonomous requirements for *Gabrb3* and *Nlgn2* in peripheral and spinal cord neuron types for tactile behaviors. **a, b**, Schematics showing areas of Cre activity (shaded in red) in spinal cord and/or dorsal root ganglia for genetic strategies shown below (top row). Adult spinal cord immunohistochemistry showing expression of VGLUT1-labeled synaptic terminals and NLGN2 in control, *Cdx2^Cre^*; *Nlgn2^flox/flox^*, and *Lbx1^Cre^*; *Nlgn2^flox/flox^*mice (**a**) and expression of VGLUT and GABRB3 in control and *Lbx1^Cre^*; *Gabrb3^flox/flox^* mice (**b**). White dotted lines show outline of spinal cord grey matter, and yellow dotted boxes and insets show dorsal horn. Scale bars denote 200 µm (full-size images) and 100 µm (insets). **c**, Tactile PPI. *P*=0.0014 for *Advillin^Cre^*; *Gabrb3^flox^*, *P*=0.6746 for *Lbx1^Cre^*; *Gabrb3^flox^*, *P*=0.0010 for *Cdx2^Cre^*; *Nlgn2^flox^*, *P*=0.7051 for *Advillin^Cre^*; *Nlgn2^flox^*, *P*<0.0001 for *Lbx1^Cre^*; *Nlgn2^flox^*, unpaired two-tailed t-tests. **d**, Response to a 0.9 PSI air puff stimulus alone. *P*=0.0011 for *Advillin^Cre^*; *Gabrb3^flox^*, *P*=0.2543 for *Lbx1^Cre^*; *Gabrb3^flox^*, *P*=0.0204 for *Cdx2^Cre^*; *Nlgn2^flox^*, *P*=0.9919 for *Advillin^Cre^*; *Nlgn2^flox^*, *P*=0.0107 for *Lbx1^Cre^*; *Nlgn2^flox^*, Mann-Whitney *U* tests. **e**, Fraction of time spent in the center of the open field chamber. *P*<0.0001 for *Advillin^Cre^*; *Gabrb3^flox^*, *P*=0.4169 for *Lbx1^Cre^*; *Gabrb3^flox^*, *P*=0.4524 for *Cdx2^Cre^*; *Nlgn2^flox^*, *P*=0.1019 for *Advillin^Cre^*; *Nlgn2^flox^*, *P*=0.4214 for *Lbx1^Cre^*; *Nlgn2^flox^*, unpaired two-tailed t-tests. **f**, Displacement response to a single presentation of 0.10, 0.25, 0.50, 0.75 and 1.0 PSI stimuli at P4. Two-way ANOVAs, effect of genotype; *Gabrb3*: (F[1, 38]=36.38, *P*<0.0001); *Nlgn2*: (F[1, 36]=0.1169, *P*=0.7344). **g**, Average displacement responses to 10 presentations of a 1.0 PSI stimulus at 20-30 second interstimulus intervals. *P*<0.0001 for *Advillin^Cre^*; *Gabrb3^flox^*, *P*=0.2382 for *Cdx2^Cre^*; *Nlgn2^flox/flox^*, unpaired one-tailed t-tests. **h**, Fraction of animals habituating to repeated presentations of 1.0 PSI air puffs. For *Gabrb3, P*=0.0171; *Nlgn2, P*=0.9333, one-sided Fisher’s Exact tests. For **c-g,** data represent means±s.e.m.

We next asked whether *Nlgn2* is required in spinal cord touch circuits for normal tactile responsiveness. We observed robust expression of NLGN2 in the spinal cord (Fig. 4a), and therefore generated *Cdx2^Cre^*; *Nlgn2^flox/flox^*^33, 54^ mice to conditionally delete *Nlgn2* in all cells caudal to cervical level 2 of the spinal cord (Fig. 4a). Unlike *Advillin^Cre^*; *Nlgn2^flox/flox^*mice, *Cdx2^Cre^*; *Nlgn2^flox/flox^* animals recapitulated the tactile behavioral deficits observed in the *Nlgn2* global knockout animals, thus revealing a novel locus of NLGN2 function outside of the brain in regulating tactile behaviors in mice (Fig. 4c,d and Extended Data Fig. 4d-f). Therefore, we turned our attention to the possibility that *Nlgn2* is required in spinal cord dorsal horn neurons, where light touch information is processed, for tactile behaviors in adult animals. To drive selective loss of *Nlgn2* from this region, we used the *Lbx1^Cre^* allele^55^, which promotes recombination in ∼95% of all neurons in the LTMR-recipient zone (LTMR-RZ) of the dorsal horn (Fig. 4a)^56^. Loss of *Nlgn2* from dorsal horn neurons in *Lbx1^Cre^*; *Nlgn2^flox/flox^* mice was sufficient to recapitulate the tactile behavioral alterations observed in both global *Nlgn2* knockout and *Cdx2^Cre^*; *Nlgn2^flox/flox^* animals (Fig. 4c,d and Extended Data Fig. 4d-f). In contrast, *Lbx1^Cre^*; *Gabrb3^flox/flox^*animals, in which the β3 subunit of GABA_A_Rs is lost from dorsal horn neurons (Fig. 4b), showed normal sensitivity to light touch stimuli (Fig. 4c,d). This may be consistent with a predominant role for glycinergic inhibition of light-touch-evoked responses in the dorsal horn^57^, and/or the presence of GABA_A_Rs that do not include the β3 subunit that are important for touch behaviors.

In keeping with our observations in adult global *Nlgn2* mutant animals, neither sensory nor spinal cord neuron loss of *Nlgn2* expression affected anxiety-like or social interaction behaviors, despite tactile over-reactivity in the spinal cord *Nlgn2* mutants (Fig. 4e and Extended Data Fig. 4g). In contrast, sensory neuron, but not spinal cord neuron, loss of *Gabrb3* expression promoted aberrant tactile, anxiety-like, and social behaviors, consistent with a specialized role for this gene in peripheral sensory neurons (Fig. 4e and Extended Data Fig. 4g). Thus, while *Gabrb3* functions cell-autonomously in peripheral somatosensory neurons, the ASD-associated gene *Nlgn2* functions in dorsal horn neurons, revealing a role for *Nlgn2* at an initial site of touch information processing in the central nervous system. Taken together with prior findings, these results indicate that both *Gabrb3* and *Mecp2* are required in peripheral somatosensory neurons^12, 22, 23^, while *Nlgn2* and *Rorb*-expressing neurons are required in the dorsal horn for tactile behaviors in adults^41^.

### Peripheral, but not spinal cord neuron locus of dysfunction predicts early emergence of tactile over-reactivity

To understand whether these distinct loci of function of the ASD-associated genes contribute to the timing differences for tactile over-reactivity during development, we next tested conditional knockout animals’ reactivity to air puff at P4. As with P4 *Gabrb3*^+/-^ animals, P4 *Advillin^Cre^*; *Gabrb3^flox/flox^* animals exhibited over-reactivity to air puff stimuli, and these conditional mutant animals also had deficits in habituation to repeated stimuli (Fig. 4f-h and Extended Data Fig. 4h, Supplementary Movie 1). Thus, the cell-autonomous requirement for *Gabrb3* in peripheral somatosensory neurons emerges early in development, and its disruption leads to developmental over-reactivity to light tactile stimuli. In contrast, and consistent with our analyses of P4 *Nlgn2*^+/-^ animals, *Cdx2^Cre^*; *Nlgn2^flox/flox^* P4 animals were indistinguishable from control littermates in their tactile reactivity (Fig. 4f-h and Extended Data Fig. 4h). Importantly, both *Advillin^Cre^*- and *Cdx2^Cre^*-mediated recombination occur during embryonic development and are complete before birth^54, 58^. Therefore, peripheral somatosensory neuron *Gabrb3* expression is required for normal tactile reactivity during early postnatal development, while spinal cord *Nlgn2* expression is dispensable during early postnatal development, but is required in adulthood.

### GABA_A_Rs that mediate presynaptic inhibition of sensory neurons are present and functional at neonatal ages

Primary somatosensory neuron-specific deletion of *Gabrb3* was sufficient to drive behavioral over-reactivity to light air puff stimuli in neonatal pups, indicating a developmental and cell-autonomous role for *Gabrb3* in peripheral sensory neurons. In adults, sensory neuron-derived GABRB3 localizes to VGLUT1^+^ sensory neuron axon terminals in the spinal cord to form GABA_A_Rs, which underlie presynaptic inhibition of these neurons^22, 23, 57^ We asked whether we would observe a similar pattern of GABRB3 expression in the neonatal spinal cord. In immunohistochemical analyses, we found that GABRB3 localized to VGLUT1^+^ axon terminals at P0-P1, and the degree of GABRB3 and VGLUT1 colocalization at this time was not significantly different from that observed in adult spinal cord tissue (Extended Data Fig. 2b,c). Furthermore, in neonatal mice, GABRB3 puncta showed synaptic apposition to VGAT^+^ presynaptic inhibitory interneuron axon terminals to a similar extent of that observed in adulthood, suggesting that the synaptic machinery underlying presynaptic inhibition of sensory afferents is in place by birth or earlier (Extended Data Fig. 2c).

To complement the histological analyses, we also asked whether peripheral sensory neurons are functionally responsive to GABA application at early postnatal ages and, if so, whether these responses require *Gabrb3* expression. To test this, we performed whole-cell patch clamp recordings of DRG neurons using an *ex vivo* preparation, and uncaged light-sensitive caged RuBi-GABA over the soma of recorded neurons (Fig. 5a). Medium- to large-diameter sensory neurons (>35 μm diameter, >55 pF capacitance)^59^, which underlie low-threshold mechanosensory responses, were targeted and dialyzed with a high-chloride internal solution. Light pulses (5 ms) consistently drove large depolarizations in these sensory neurons at P4, and their magnitudes were not significantly different from P18–30 neuron responses (Fig. 5b,c). Furthermore, at both timepoints, GABA-evoked responses in medium- and large-diameter DRG neurons of *Advillin*^Cre^; *Gabrb3^flox/flox^* animals were virtually abolished, highlighting the obligatory role of GABRB3 for the formation of functional GABA_A_Rs in these peripheral sensory neurons (Fig. 5b,c). Thus, GABA_A_Rs are present on medium- and large-diameter DRG neurons during neonatal development, and require *Gabrb3* expression for responses to GABA. Interestingly, although presynaptic GABA_A_Rs are present early in development, top-down cortical inputs into the dorsal horn, which engage presynaptic inhibition of primary afferents and other aspects of local touch processing in adulthood^56, 60^, were anatomically absent (Extended Data Fig. 5a). Together, our immunohistochemical and electrophysiological analyses indicate that the synaptic machinery underlying presynaptic inhibition of primary somatosensory neuron afferents has matured by neonatal ages, supporting the idea that ASD-associated mutations impacting presynaptic inhibition disrupt sensory neuron function and tactile reactivity very early in development.

**Figure 5.**
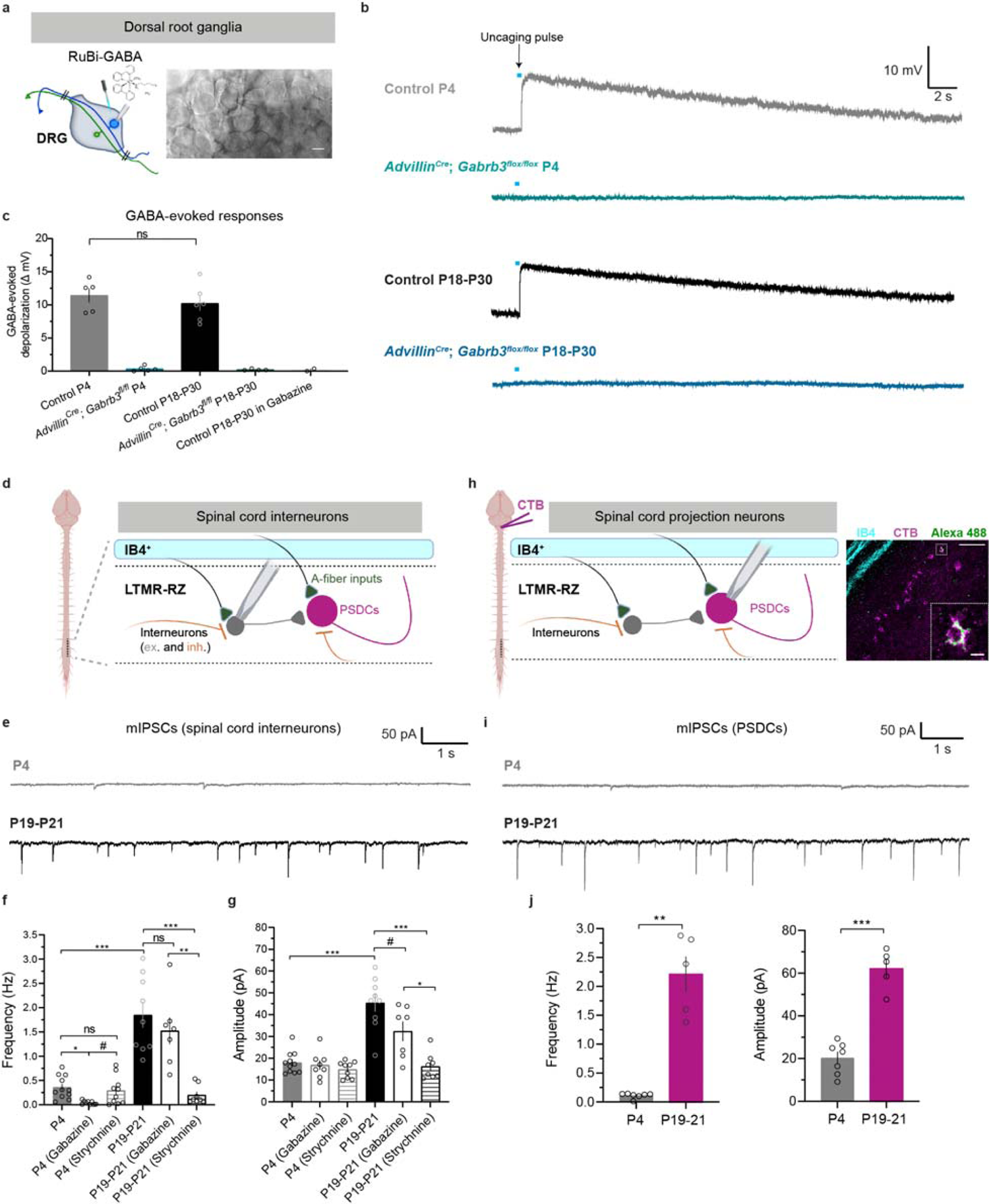
Machinery for presynaptic inhibition of sensory neurons develops neonatally, while spinal cord feedforward inhibition is weak and immature during early postnatal development. **a**, Experimental setup and example image of DRG whole-cell recording preparation for RuBi-GABA uncaging. Scale bar denotes 25 µm. **b**, Example GABA-evoked depolarizations of medium- to large-diameter neurons to a 5 ms uncaging pulse (blue bar, not to scale) in current-clamp configuration. **c**, Quantification of peak uncaged GABA-evoked depolarizations across time points and genotypes. Each dot represents a cell, and N=2 animals for each condition, except N=3 for Control P18-30. *P*=0.4592, unpaired two-tailed t-test. **d**, Recording schematic for measuring miniature postsynaptic currents from interneurons (both excitatory and inhibitory, denoted ex. and inh.) in the low-threshold mechanoreceptor recipient zone (LTMR-RZ, laminae III-IV, which resides underneath the IB4^+^ superficial lamina II) of sagittal spinal cord slices. **e**, Example miniature inhibitory post-synaptic currents (mIPSCs) in P4 and P19-21 spinal cords. Recordings were performed in lamina III/IV of sagittal spinal cord sections. **f**, **g**, mIPSC frequencies (**f**) and amplitudes (**g**) at P4 and P19-21, with or without the presence of blockers of synaptic inhibition, gabazine and strychnine. N=3 animals for each condition, except N=2 for P4 (gabazine), P4 (strychnine), and P19-21 (strychnine). For frequencies, *P*=0.0105 for P4 vs. P4 (gabazine), *P*=0.7714 for P4 vs. P4 (strychnine), *P*=0.0550 for P4 (gabazine) vs. P4 (strychnine), *P*<0.0001 for P4 vs. P19-P21, *P*=0.5701 for P19-P21 vs. P19-P21 (gabazine), *P*<0.0001 for P19-P21 vs. P19-P21 (strychnine), and *P*=0.0015 for P19-P21 (gabazine) vs. P19-P21 (strychnine). For amplitudes, *P*=0.8821 for P4 vs. P4 (gabazine), *P*=0.3553 for P4 vs. P4 (strychnine), *P*=0.6827 for P4 (gabazine) vs. P4 (strychnine), *P*<0.0001 for P4 vs. P19-P21, *P*=0.0521 for P19-P21 vs. P19-P21 (gabazine), *P*<0.0001 for P19-P21 vs. P19-P21 (strychnine), and *P*=0.0173 for P19-P21 (gabazine) vs. P19-P21 (strychnine). One-way ANOVAs with post hoc Tukey’s tests. **h**, CTB was injected into the dorsal column or dorsal column nuclei to retrogradely label postsynaptic dorsal column neurons (PSDCs). Sagittal spinal cord slices were prepared and PSDCs were targeted for whole-cell patch clamp recordings (left), and in some cases, dialyzed with Alexa Fluor 488 dye for post hoc visualization (right). Scale bars denote 100 µm (full-size image) and 10 µm (inset). **i**, Example miniature inhibitory post-synaptic currents (mIPSCs) from CTB-labeled PSDCs in P4 and P19-21 spinal cords. **j**, mIPSC frequencies (left) and amplitudes (right) at P4 and P19-21 onto PSDCs. N=3 animals for both conditions. For frequencies, *P*=0.0024, unpaired two-tailed Welch’s t-test. For amplitudes, *P*<0.0001, unpaired t test. For **c**, **f**, **g**, and **j**, data represent means±s.e.m.

### Feedforward inhibition of spinal cord neurons is immature at early postnatal stages

In contrast to *Gabrb3* and *Mecp2* mutant animals, *Nlgn2* and *Rorb* mutant animals exhibited normal tactile behaviors during early postnatal development. However, spinal cord-specific disruptions of *Nlgn2* and *Rorb* neurons drive aberrant reactivity in adulthood (Fig. 4c,d; ref ^41^). We therefore hypothesized that the spinal cord functions mediated by *Nlgn2* and *Rorb* develop after early postnatal periods, in contrast to the early developmental sensory neuron functions of *Gabrb3* and *Mecp2*^23^. Since both *Nlgn2* and *Rorb* are implicated in feedforward inhibitory neurotransmission^31, 41, 61^, we asked whether feedforward inhibition of dorsal horn LTMR-RZ interneurons and projection neurons (lamina III-IV of the spinal cord) was present at early neonatal stages. Therefore, we measured miniature inhibitory post-synaptic currents in lamina III-IV of neonatal (P4) and mature (P19-P21) animals (Fig. 5d). At P4, the frequency and amplitude of mIPSCs onto spinal cord neurons were dramatically lower compared to those of P19-P21 spinal cord neurons (Fig. 5e-g). Furthermore, while application of the GABA_A_R antagonist gabazine blocked nearly all mIPSCs at P4, gabazine did not strongly affect mIPSC frequency in recordings from P19-P21 mice (Fig. 5f). Application of the glycine receptor antagonist strychnine reduced both the frequency and amplitude of inhibitory events in mature slices, but not at P4 (Fig. 5f,g). In contrast, AMPA receptor-mediated mEPSCs in spinal cord neurons were more similar early and late in development (Extended Data Fig. 5b).

In complementary experiments, we recorded mIPSCs from retrogradely-labeled postsynaptic dorsal column neurons (PSDCs), which are dorsal horn output neurons that project to the brainstem^62^ and receive direct inputs from Aβ LTMRs, other LTMR subtypes, and spinal cord interneurons^41, 56^ (Fig. 5h). Consistent with randomly recorded lamina III-IV neurons, the frequency and amplitude of mIPSCs in PSDCs from P4 pups were markedly lower than those of mature PSDCs from P19-P21 mice (Fig. 5i, j).

Finally, measurements of VGAT^+^ inhibitory interneuron axon terminals and post-synaptic alpha 1 subunit-containing glycine receptors in the spinal cord revealed that synaptic apposition between VGAT and glycine receptors was significantly lower at P4 compared to that of adult spinal cord tissue^57^ (Extended Data Fig. 5c,d). Together, these findings indicate that feedforward inhibition in the LTMR-RZ develops late: inhibition is weak and mediated solely by GABA at P4, and it greatly strengthens and becomes predominantly glycinergic by P19. These findings are consistent with prior analyses performed in the superficial dorsal horn of developing rat spinal cords, where nociceptive inputs impinge on spinal neurons characterized by a lack of inhibitory control and a late emergence of glycinergic inhibition^63^.

### Neuroligin-2 is required for feedforward inhibitory neurotransmission in the mature spinal cord dorsal horn

Our behavioral findings demonstrate that NLGN2 is required for normal tactile reactivity in adulthood but not at P4, and our electrophysiological recordings indicate that feed-forward inhibitory synaptic transmission in the LTMR-RZ is robust in adult dorsal horn but nearly absent at P4. Thus, we hypothesized that NLGN2 functions during late postnatal development of the dorsal horn LTMR-RZ to control the maturation of inhibitory transmission. In spinal cord slice preparations, mIPSCs recordings in the LTMR-RZ revealed that both the frequency and amplitude of mIPSCs were markedly reduced in P19-21 *Cdx2^Cr^*^e^; *Nlgn2^flox/flox^* slices compared to controls (Fig. 6a,b). Consistent with previous work in other central nervous system regions^30, 33, 64–66^, this change was specific to inhibitory neurotransmission: mEPSC frequency and amplitude were normal in the same preparation (Fig. 6c). Thus, in late postnatal mice, *Nlgn2* is required for feedforward inhibitory neurotransmission in the spinal cord dorsal horn, where light-touch signals are first relayed. In contrast, and consistent with our behavioral findings, recordings in P4–P5 *Cdx2^Cre^*; *Nlgn2^flox/flox^* and littermate control spinal cords revealed no significant differences in mIPSC frequency or amplitude between groups (Fig. 6d). Finally, immunohistochemical analyses of the expression of both NLGN2 and gephyrin, an inhibitory postsynaptic scaffolding gene, in the neonatal spinal cord showed no detectable signal at P4, despite robust expression in adulthood (Extended Data Fig. 6a-c). These electrophysiological and histological analyses indicate that NLGN2 is lowly or not expressed and dispensable for the weak and immature feedforward inhibition present in the early postnatal deep dorsal horn, consistent with the observation that *Nlgn2* deletion does not affect tactile reactivity during early postnatal development. In contrast, by late postnatal ages and adulthood, NLGN2 is robustly expressed and required for normal feedforward inhibitory transmission in the spinal cord, which is primarily glycinergic. Thus, the cellular and synaptic mechanisms underlying glycinergic feedforward inhibition and GABAergic presynaptic inhibition in the spinal cord mature at different stages. These findings support a model in which ASD-associated gene mutationsaffecting presynaptic inhibition of LTMRs manifest during late embryonic and early neonatal development, and thereby have profound effects on embryonic and early postnatal reactivity to tactile stimuli. In contrast, ASD-associated gene mutations affecting dorsal horn feedforward inhibition manifest later, during late postnatal development, and thus are without deleterious consequence in early postnatal tactile reactivity (Fig. 6e and Extended Data Fig. 6d). Importantly, animals with tactile over-reactivity neonatally also exhibit increased anxiety-like behaviors and decreased sociability in adulthood, whereas animals with normal early tactile reactivity do not exhibit these changes.

**Figure 6:**
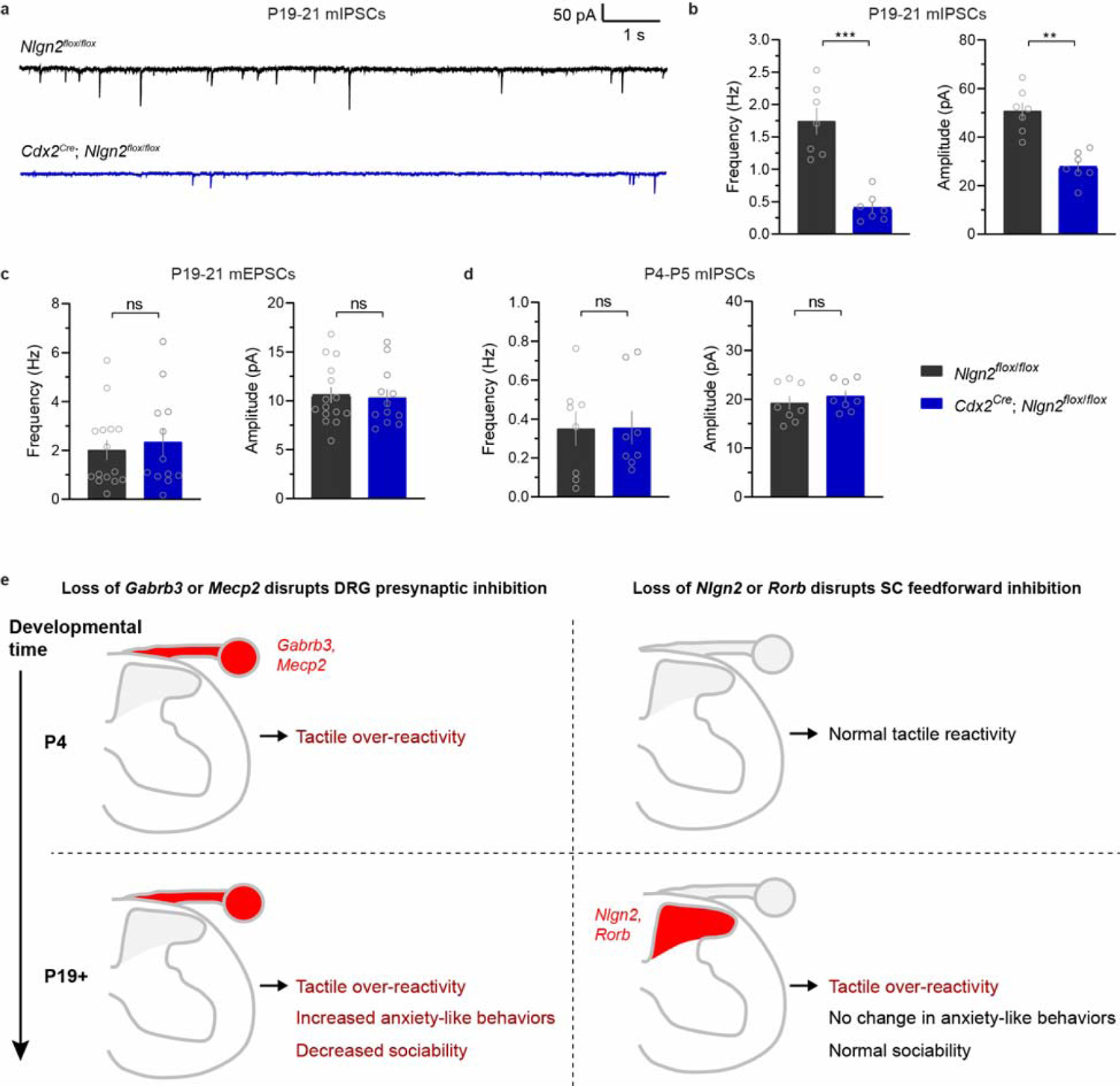
Neuroligin-2 is required for feedforward inhibitory neurotransmission in the mature spinal cord dorsal horn. **a**, Example mIPSCs in spinal cord neurons of P19-21 spinal cords in *Cdx2^Cre^*; *Nlgn2^flox/flox^* animals and control littermates. **b**, mIPSC frequencies (left) and amplitudes (right) of P19-21 spinal cords in *Cdx2^Cre^*; *Nlgn2^flox/flox^* animals and control littermates. N=2 animals for both conditions. *P*<0.0001 for frequencies, and *P*=0.0001 for amplitudes, unpaired two-tailed t-tests. **c**, Frequencies (left) and amplitudes (right) of mEPSCs in the spinal cords of *Cdx2^Cre^*; *Nlgn2^flox/flox^* animals and control littermates at P19-21. N=2 animals for both conditions. *P*=0.6384 for frequencies, and *P*=0.6822 for amplitudes, unpaired two-tailed t-tests. **d**, Frequencies (left) and amplitudes (right) of mIPSCs in the spinal cords of *Cdx2^Cre^*; *Nlgn2^flox/flox^* animals and control littermates at P4-P5. N=2 animals for both conditions. *P*=0.9678 for frequencies, and P=0.4385 for amplitudes, unpaired two-tailed t-tests. **e**, Proposed model describing the emergence of dorsal root ganglia (DRG) presynaptic inhibition and spinal cord (SC) feedforward inhibition across development, with observed behavioral consequences of disrupting ASD-associated genes involved in either pathway.

## DISCUSSION

Here, we report that distinct cellular and synaptic mechanisms in peripheral somatosensory neurons and spinal cord neurons contribute to tactile over-reactivity in mouse models of ASDs. Mice with deletion of the ASD-associated genes *Gabrb3* or *Mecp2*, which are required for presynaptic inhibition onto peripheral somatosensory afferents^12, 22, 23^, exhibit aberrant tactile reactivity from early neonatal stages. In contrast, disruptions in *Nlgn2* or *Rorb*, which impact feedforward inhibition of spinal cord dorsal horn neurons^41^, do not affect tactile reactivity neonatally. While all of these animal models exhibit aberrant touch behaviors in adulthood, mice that had tactile over-reactivity in early life also present with anxiety-like behaviors and sociability deficits in adulthood, whereas mice with normal tactile reactivity in early life do not exhibit these additional alterations. Therefore, in these models of ASD, developmental somatosensory dysfunction is linked to the emergence of cognitive and social behavior disruptions.

We found that the machinery for GABAergic presynaptic inhibition of large-diameter peripheral somatosensory neurons is present by early neonatal stages. Therefore, ASD-associated mutations that disrupt presynaptic GABA_A_ receptors impact somatosensory behaviors during development. To assess this, we developed an approach to quantitatively measure tactile reactivity in neonatal animals, and found that the *Gabrb3* and *Mecp2* loss-of-function models exhibited tactile over-reactivity to a gentle air puff stimulus at P4. This over-reactivity can be attributed to a cell-autonomous role for GABAergic signaling in peripheral somatosensory neurons: somatosensory neuron-specific deletion of *Gabrb3* ablated functional responsivity to GABA in sensory neurons, and mutant animals recapitulated the aberrant behaviors we observed in *Gabrb3*^+/-^ mice. Remarkably, *Gabrb3*^+/-^ mice also exhibited over-reactivity to activation of Ret^+^ Aβ LTMRs, during late embryonic development, and so loss of primary afferent presynaptic inhibition may drive sensory over-reactivity *in utero*. Consistent with this, a prior study detected GABAergic dorsal root potentials, a hallmark of presynaptic inhibition, as early as E17.5 in isolated rat spinal cords^67^. We therefore speculate that aberrant activity in tactile circuits *in utero* or early postnatal development may contribute to the emergence of core and co-morbid ASD features in certain individuals. This finding extends our prior work defining a critical period that ends during early postnatal life in mice, in which physiological dysfunction of primary sensory neurons impacted cortical microcircuits and drove enhanced anxiety-like behaviors and abnormal social interactions^12^.

While the machinery that underlies presynaptic inhibition of peripheral sensory neurons matures early in development, feedforward inhibitory neurotransmission onto central neurons in the spinal cord dorsal horn, where light touch is processed, matures across a much slower timescale. Our work in the mouse low-threshold mechanoreceptor recipient zone (LTMR-RZ) agrees with studies performed in the rat superficial spinal cord, showing that feedforward inhibition onto spinal cord neurons shifts from early GABAergic inputs to glycinergic control over the course of postnatal development^63, 68, 69^. In the adult rat, LTMR-evoked responses are under strong glycinergic control^57, 70^, and it is thought that both glycinergic inputs and postsynaptic receptors only reach adult levels of maturity at the end of the second postnatal week^57, 69^. Here, we identified *Nlgn2* as an important component in feedforward inhibition of dorsal horn neurons. Developmental deletion of *Nlgn2* did not impact spinal synaptic function or behavioral reactivity to low-threshold stimuli during early postnatal development, and NLGN2, the inhibitory postsynaptic scaffolding protein gephyrin, and glycine receptor expression were undetectable in the early postnatal spinal cord. The emergence of aberrant tactile behaviors at late postnatal times in *Nlgn2* mutants likely corresponds to the late impact on the maturation of feedforward inhibitory circuitry. In related experiments, we investigated the ASD-associated gene *Rorb*, because silencing *Rorb*-expressing spinal neurons leads to a dramatic loss of glycinergic feedforward inhibition onto PSDCs and promotes tactile dysfunction^41^. Similar to *Nlgn2* mutant animals, *Rorb* gene mutant animals had normal touch reactivity at P4 but exhibited augmented tactile reactivity in adulthood. There was substantial correspondence in behavioral phenotypes between animals with *Rorb* gene mutations versus silenced or ablated *Rorb* neurons^41^, but future studies should address whether spinal cord feedforward synaptic inhibition is impacted to the same extent in these two conditions. Furthermore, while deficits in spontaneous post-synaptic currents were observed in *Nlgn2* mutant animals in the present study, it remains to be determined whether touch-evoked spiking activity is altered in the spinal cords of these *Nlgn2* mutants during postnatal development or adulthood^71^.

Our findings support a model in which NLGN2 is an essential organizer of glycinergic synapses^30^ that lie postsynaptic to inhibitory glycinergic *Rorb*-expressing and other inhibitory interneurons of the deep dorsal horn (Extended Data Fig. 6d). Thus, *Nlgn2* and *Rorb* may function within a common feedforward inhibition pathway in the mature dorsal horn that regulates behavioral responses to touch *after* a critical time window of perinatal development, during which tactile signals shape the central circuits governing non-tactile ASD co-morbidities^12^. Indeed, *Nlgn2* and *Rorb* mutant animals exhibit normal anxiety-like and social behaviors in adulthood, although it is possible that these animals may exhibit behavioral alterations not examined here or in prior studies^35, 44^. Taken together, we propose that the differences in the developmental timing of function of ASD-associated genes that act within the same circuitry, for example, *Gabrb3*, *Mecp2*, *Nlgn2*, and *Rorb* in the DRG sensory neuron/dorsal horn circuitry, help to explain the heterogeneity of behavioral outcomes observed across ASD mouse models.

In humans with ASD, behavioral studies indicate that altered sensory behaviors often predict the severity of higher-order ASD or ASD-related traits^8, 14, 72–76^. Importantly, altered sensory reactivity can be detected during early development in subsets of children with ASD^77^, suggesting that sensory processing changes may represent an early clinically relevant marker for the emergence of future ASD-related traits in subgroups of individuals. Indeed, the concept of “chronogeneity” describes the diverse clinical trajectories that individuals with ASD can take through development and onward, and posits that temporal and cross-sectional heterogeneity across individuals may represent a form of “informative variance” that describes a more dynamic and well-powered understanding of ASDs^78^. Our findings lead us to propose that variability in spinal sensory circuit alterations in ASD conditions, and therefore variable developmental timing of aberrant sensory behaviors, contributes to observed chronogeneity across individuals. Therefore, a richer understanding of how atypical sensory processing behaviors during early development relate to other core and co-morbid features of ASDs may inform future diagnostic, prognostic, and interventional strategies.

## EXTENDED DATA FIGURES

**Extended Data Figure 1.**
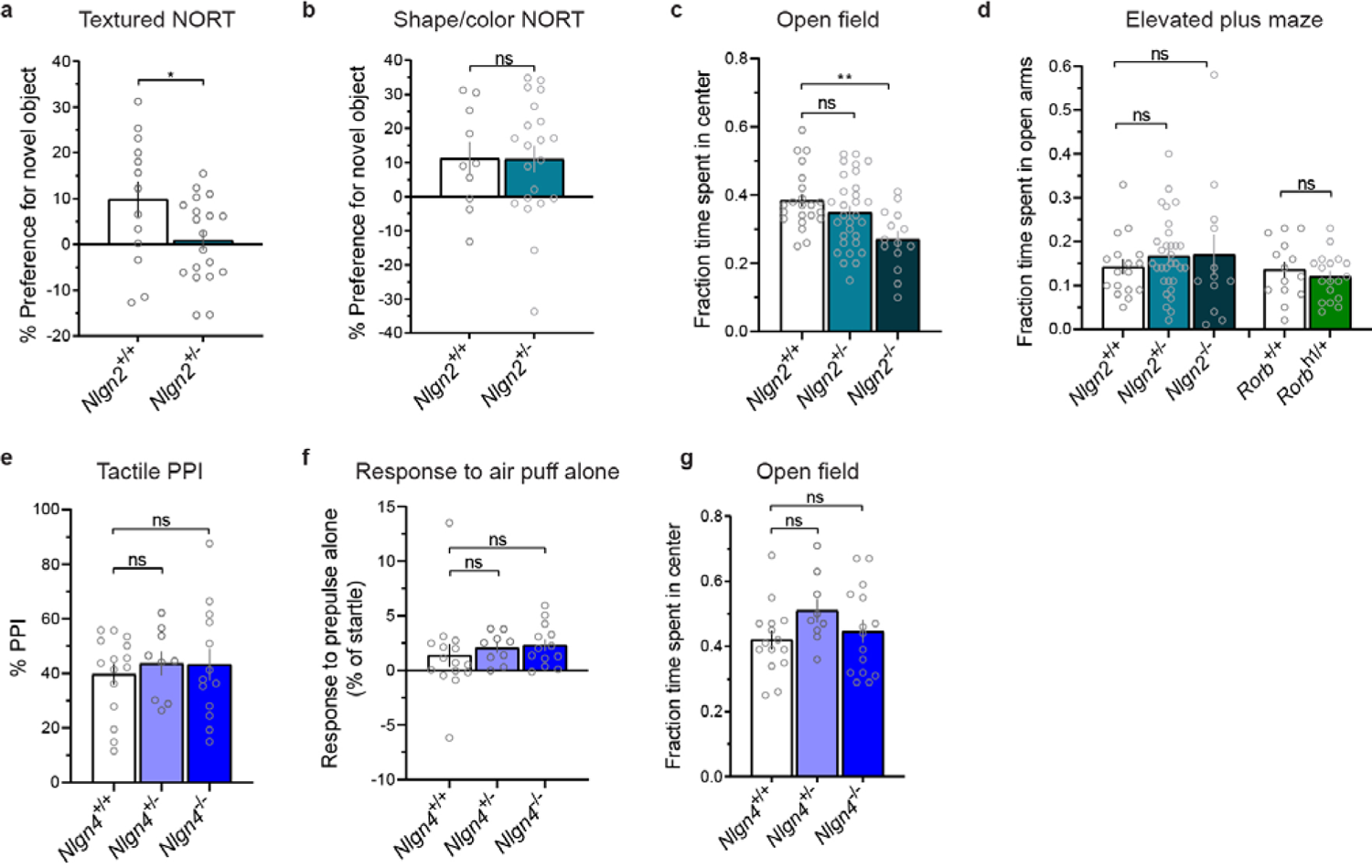
Additional behavioral characterization of global knockout ASD models. Related to **Fig. 1**. **a, b**, Discrimination indices for textured NORT, *P*LJ=LJ0.0376 (**a**) and shape/color NORT, *P*LJ=LJ0.9729 (**b**) assays, with a positive value indicating a preference for a novel object versus a familiar object. Statistical analyses were performed using unpaired two-tailed t-tests. **c**, Fraction of time spent in the center of the open field chamber. *P*=0.3289 for *Nlgn2*^+/+^ vs. *Nlgn2*^+/-^, *P*=0.0020 for *Nlgn2*^+/+^ vs. *Nlgn2*^-/-^, one-way ANOVA with post hoc Dunnett’s test. **d**, Fraction of time spent in the open arms of the elevated plus maze. *P*=0.6302 for *Nlgn2*^+/+^ vs. *Nlgn2*^+/-^, *P*=0.6886 for *Nlgn2*^+/+^ vs. *Nlgn2*^-/-^, one-way ANOVA with post hoc Dunnett’s test, and *P*=0.4630 for *Rorb*, unpaired two-tailed t-tests. **e**, Tactile PPI. *P*=0.2674 for *Nlgn4*^+/+^ vs. *Nlgn4*^+/-^, *P*=0.1912 for *Nlgn4*^+/+^ vs. *Nlgn4*^-/-^, Kruskal-Wallis test with post hoc Dunn’s test. **f**, Response to a 0.9 PSI air puff stimulus alone. *P*=0.8100 for *Nlgn4*^+/+^ vs. *Nlgn4*^+/-^, *P*=0.6223 for *Nlgn4*^+/+^ vs. *Nlgn4*^-/-^. **g**, Fraction of time spent in the center of the open field chamber. *P*=0.1461 for *Nlgn4*^+/+^ vs. *Nlgn4*^+/-^, *P*=0.7818 for *Nlgn4*^+/+^ vs. *Nlgn4*^-/-^. For **a-g**, data represent means±s.e.m. For **f** and **g**, statistical analyses were performed using one-way ANOVA with post hoc Dunnett’s tests.

**Extended Data Figure 2.**
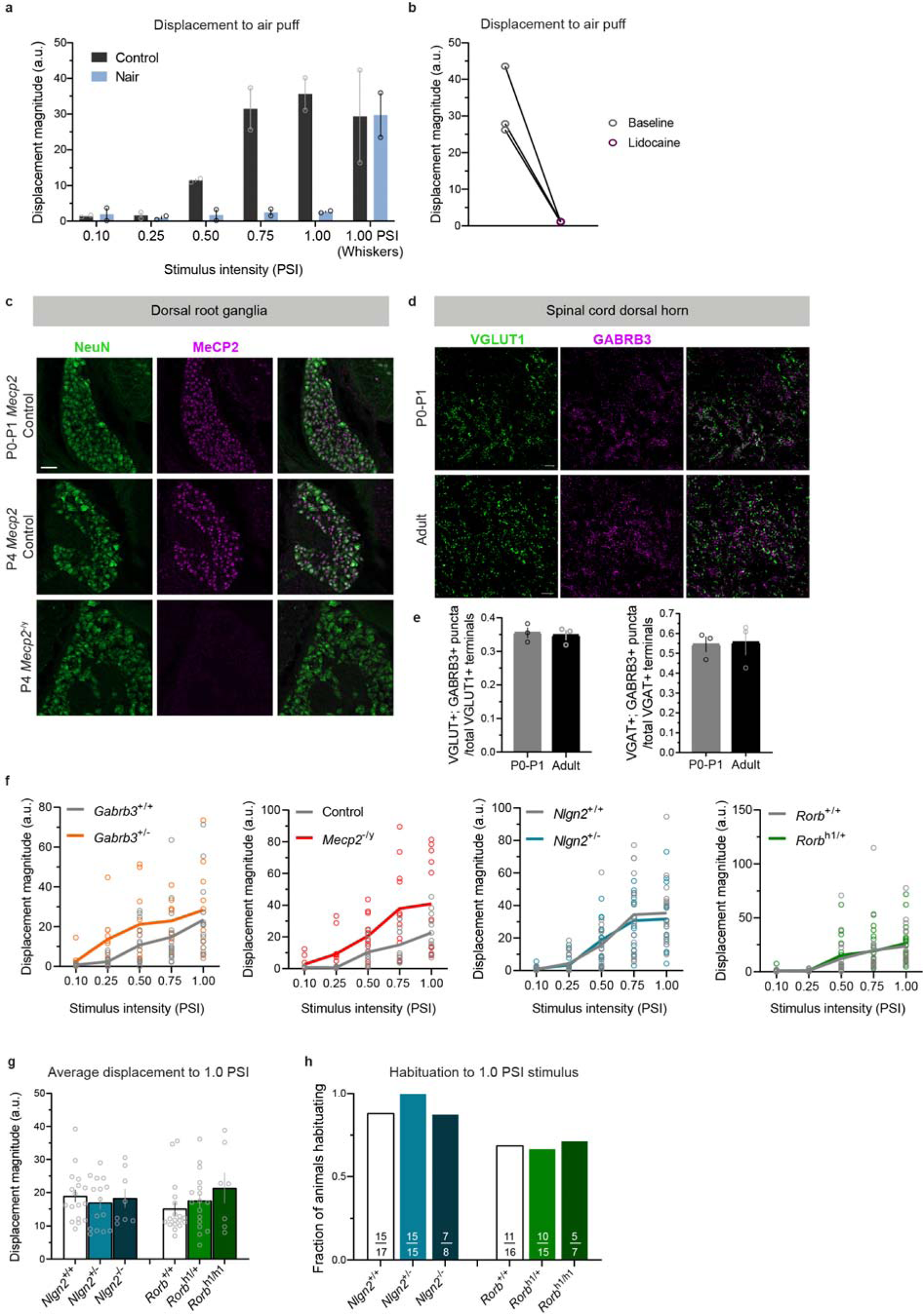
Additional behavioral and molecular characterization of ASD mouse models during neonatal ages. Related to **Fig. 2**. **a**, Displacement responses to single air puff stimuli of varying intensities delivered in sequence to the nape of the neck, in P4 animals treated with Nair on the back hairy skin and non-treated animals. At the end of back hairy skin trials, the 1.0 PSI stimulus was delivered to the whisker pad. **b**, Displacement responses to a single 1.0 PSI air puff stimulus before and after topical lidocaine application to the nape of the neck. Each pair of dots represents the same P4 animal pre- and post-application. **c**, Dorsal root ganglia immunohistochemistry showing expression of NeuN and MeCP2 in control mice at P0-P1 and P4, and in *Mecp2*^-/y^ mice to validate antibody specificity. Scale bar denotes 50 µm. **d**, Spinal cord immunohistochemistry showing expression of VGLUT1-labeled synaptic terminals and GABRB3 in control P0-P1 and adult mice. Scale bars denote 10 µm. **e**, Quantification of VGLUT1 and GABRB3 (left) or VGAT and GABRB3 (right) co-labeled puncta, as a fraction of total VGLUT1+ or VGAT+ puncta in a field of view, at P0-P1 or adulthood. Each dot represents the average value for one animal, with 3 spinal cord images taken per animal. **f**, Displacement responses to a single presentation of 0.10, 0.25, 0.50, 0.75 and 1.0 PSI stimuli for global mutant animals. Each dot represents the displacement response of a single animal on an air puff trial. **g**, Average displacement responses to 10 presentations of a 1.0 PSI stimulus at 20-30 second interstimulus intervals. *P*=0.6848 for *Nlgn2*^+/+^ vs. *Nlgn2*^+/-^, *P*=0.9749 for *Nlgn2*^+/+^ vs. *Nlgn2*^-/-^, and *P*=0.6596 for *Rorb*^+/+^ vs. *Rorb*^h1/+^, *P*=0.2211 for *Rorb*^+/+^ vs. *Rorb*^h1/h1^, one-way ANOVA with post hoc Dunnett’s tests. **h**, Fraction of animals habituating to repeated presentations of 1.0 PSI air puffs. For **a**, **e,** and **g**, data represent means±s.e.m.

**Extended Data Figure 3.**
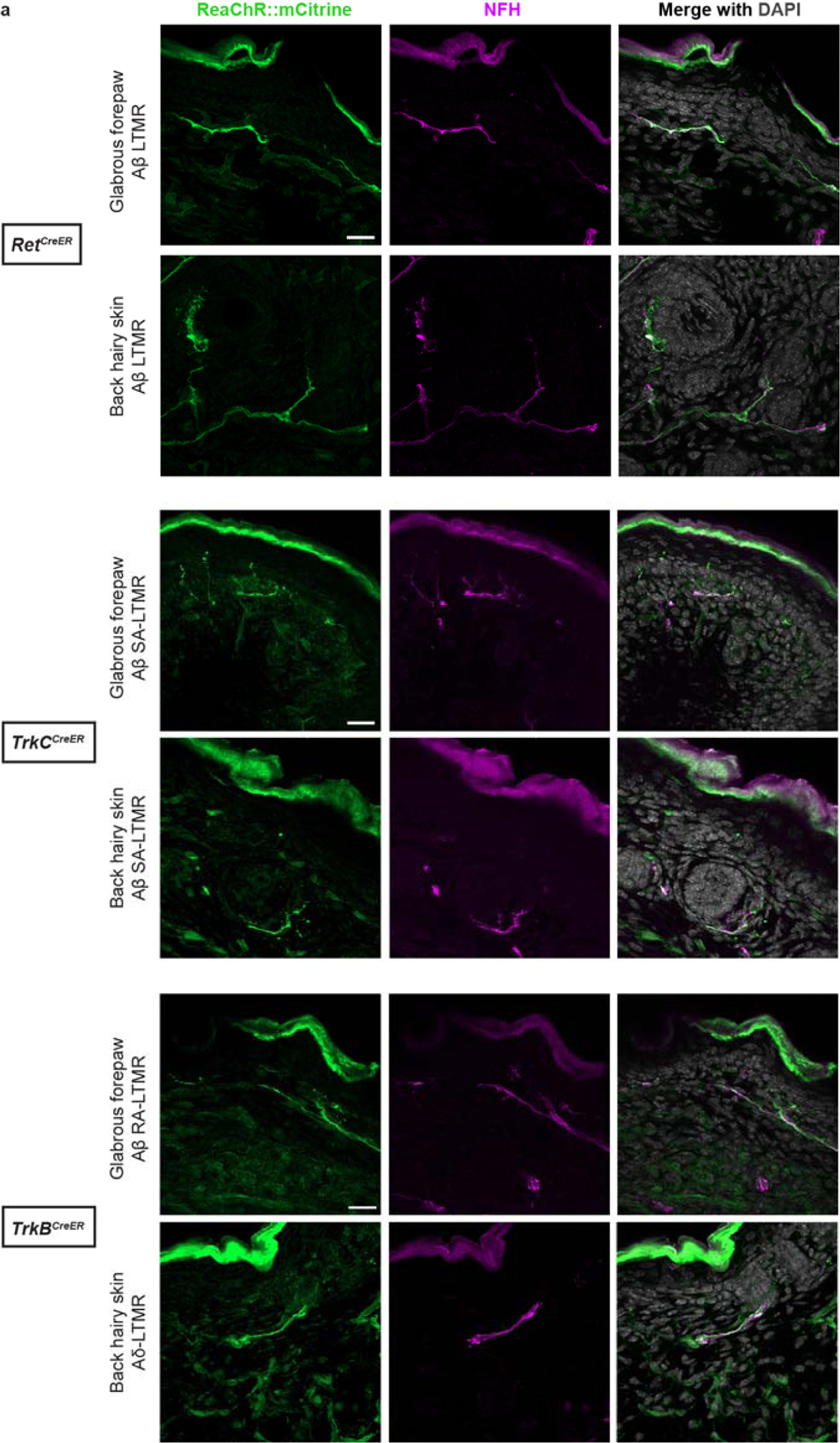
Anatomical characterization of opsin expression in low-threshold mechanoreceptor labeling paradigm at birth. Related to **Fig. 3**. **a**, Glabrous forepaw and back hairy skin immunohistochemistry showing expression of ReaChR::mCitrine (green), Neurofilament Heavy Chain (NFH; magenta), and DAPI (gray) from P0 animals with *Ret^CreER^*, *TrkC^CreER^*, or *TrkB^CreER^* mechanosensory neuron labeling strategies. Scale bars denote 20 µm.

**Extended Data Figure 4.**
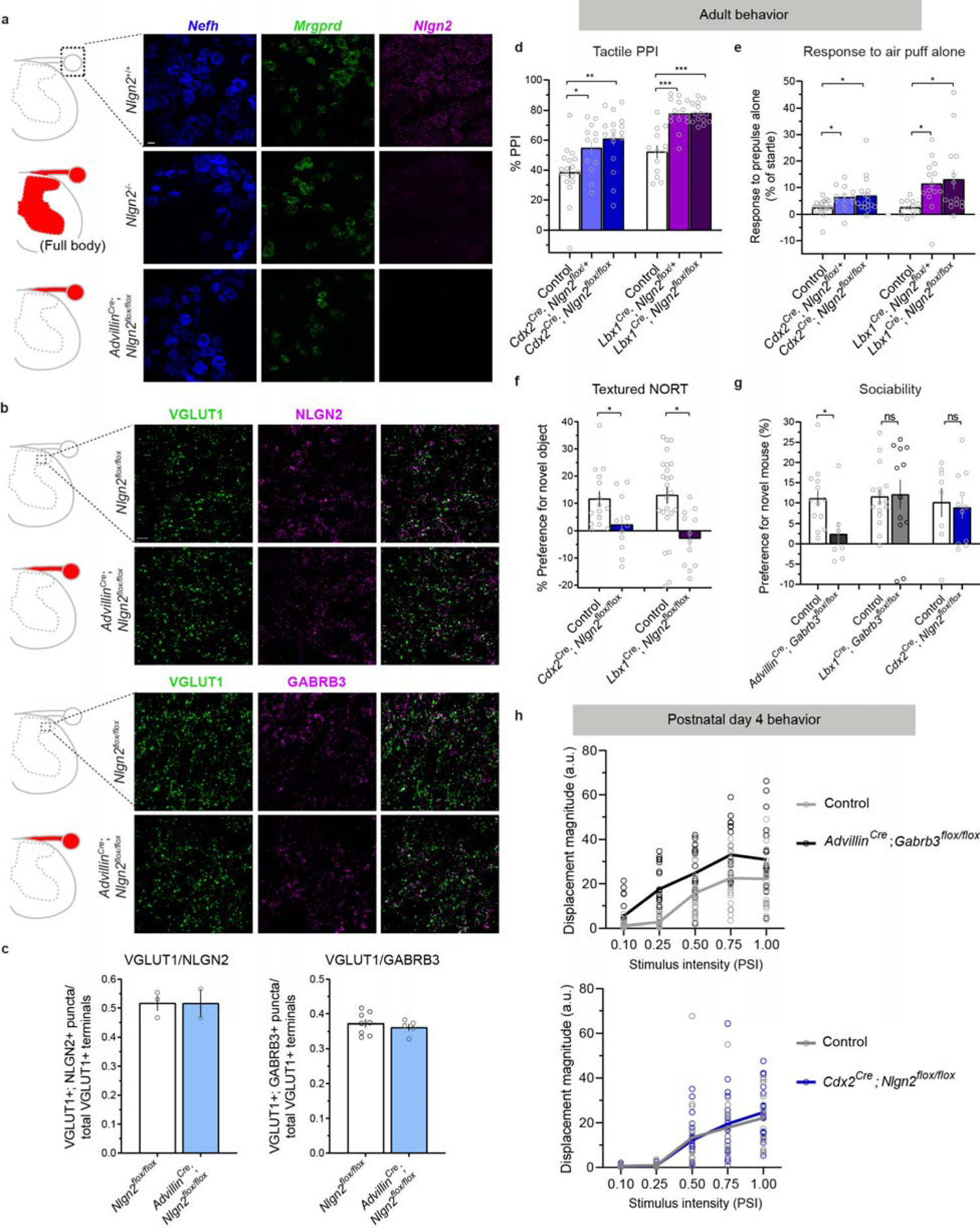
Additional characterization of conditional knockout ASD models. Related to **Fig. 4**. **a**, Schematics showing imaging area (black dotted box) and areas of Cre activity, where relevant (shaded in red) (left). *In situ* hybridization labeling using RNAScope for *Nefh* (large-diameter DRG neurons), *Mrgprd* (small-diameter nonpeptidergic DRG neurons), and *Nlgn2* transcripts in DRG cells of *Nlgn2*^+/+^, *Nlgn2*^+/-^, and *Advillin^Cre^*; *Nlgn2^flox/flox^* mice (right). Scale bar denotes 20 µm. **b**, Schematics showing imaging area (black dotted box) and areas of Cre activity, where relevant (shaded in red) (left). Spinal cord immunohistochemistry showing expression of VGLUT1-labeled synaptic terminals and NLGN2 (top, right) and GABRB3 (bottom, right) in control and *Advillin^Cre^*; *Nlgn2^flox/flox^* mice. Scale bar denotes 10 µm. **c**, Quantification of VGLUT1 and NLGN2 (left) or VGLUT1 and GABRB3 (right) co-labeled puncta, as a fraction of total VGLUT1+ puncta in a field of view. Each dot represents the average value for one animal, with 3 spinal cord images taken per animal. **d**, Tactile PPI. *P*=0.0367 for Control vs. *Cdx2^Cre^*; *Nlgn2^flox/+^*, *P*=0.0012 for Control vs. *Cdx2^Cre^*; *Nlgn2^flox/flox^*, *P*<0.0001 for Control vs. *Lbx1^Cre^*; *Nlgn2^flox/+^*, *P*<0.0001 for Control vs. *Lbx1^Cre^*; *Nlgn2^flox/flox^*, one-way ANOVA with post hoc Dunnett’s tests. **e**, Response to a 0.9 PSI air puff stimulus alone. *P*=0.0181 for Control vs. *Cdx2^Cre^*; *Nlgn2^flox/+^*, *P*=0.0383 for Control vs. *Cdx2^Cre^*; *Nlgn2^flox/flox^*, *P*=0.0014 for Control vs. *Lbx1^Cre^; Nlgn2^flox/+^*, *P*=0.0174 for Control vs. *Lbx1^Cre^*; *Nlgn2^flox/flox^*, Kruskal-Wallis tests with post hoc Dunn’s tests. **f**, Discrimination indices for textured NORT. *P=*0.0357 for *Cdx2^Cre^*; *Nlgn2^flox^*, *P=*0.0028 for *Lbx1^Cre^*; *Nlgn2^flox^*, unpaired two-tailed t-tests. **g**, Percent preference for a novel mouse over a novel object in the 3-chamber social interaction assay. *P*=0.0217 for *Advillin^Cre^*; *Gabrb3^flox^*, *P*=0.8816 for *Lbx1^Cre^*; *Gabrb3^flox^*, *P*=0.7776 for *Cdx2^Cre^*; *Nlgn2^flox^*, unpaired two-tailed t-tests. **h**, Displacement responses to a single presentation of 0.10, 0.25, 0.50, 0.75 and 1.0 PSI stimuli for P4 *Advillin^Cre^*; *Gabrb3^flox/flox^* (top) and *Cdx2^Cre^*; *Nlgn2^flox/flox^* (bottom) and control littermate animals. Each dot represents the displacement response of a single animal on an air puff trial. For **c**-**g**, data represent means±s.e.m.

**Extended Data Figure 5.**
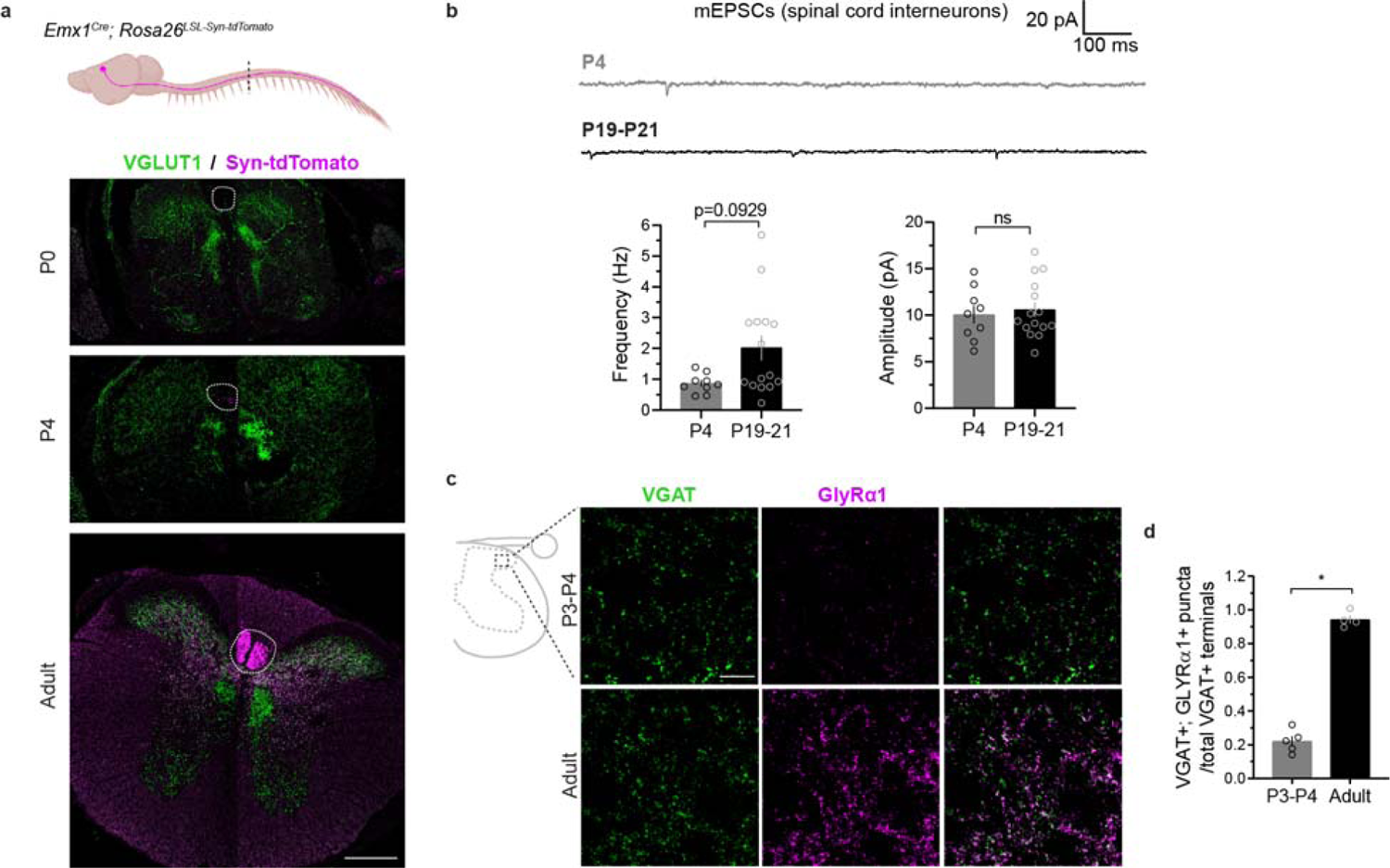
Synaptic elements for spinal cord inhibition emerge at different times, and additional characterization of neurotransmission in developing spinal cords. Related to **Fig 5**. a, Anatomical labeling of the corticospinal tract in thoracic spinal cords of P0, P4, and adult animals labeled by in *Emx1^Cre^*; *Rosa26^LSL-synaptophysin-tdTomato^*. Scale bar denotes 200 µm. Parts created with <u>BioRender.com</u>. b, Example miniature excitatory post-synaptic currents (mEPSCs) from P4 and P19-21 spinal cord interneurons of lamina III/IV (top), and quantification of frequencies (bottom, left) and amplitudes (bottom, right). N=2 animals for P4 and N=3 for P19-21. For frequencies, *P*=0.0929, Mann-Whitney *U* test. For amplitudes, *P*=0.6822, unpaired two-tailed t-test. c, Spinal cord immunohistochemistry showing expression of VGAT-labeled synaptic terminals and GLYRα1 in P3-P4 and adult mice. Scale bar denotes 10 µm. d, Quantification of VGAT and GLYRα1 co-labeled puncta, as a fraction of total VGAT+ puncta in a field of view, at P3-P4 or adulthood. *P*<0.0001, unpaired two-tailed t-test. For b and d, data represent means±s.e.m.

**Extended Data Figure 6.**
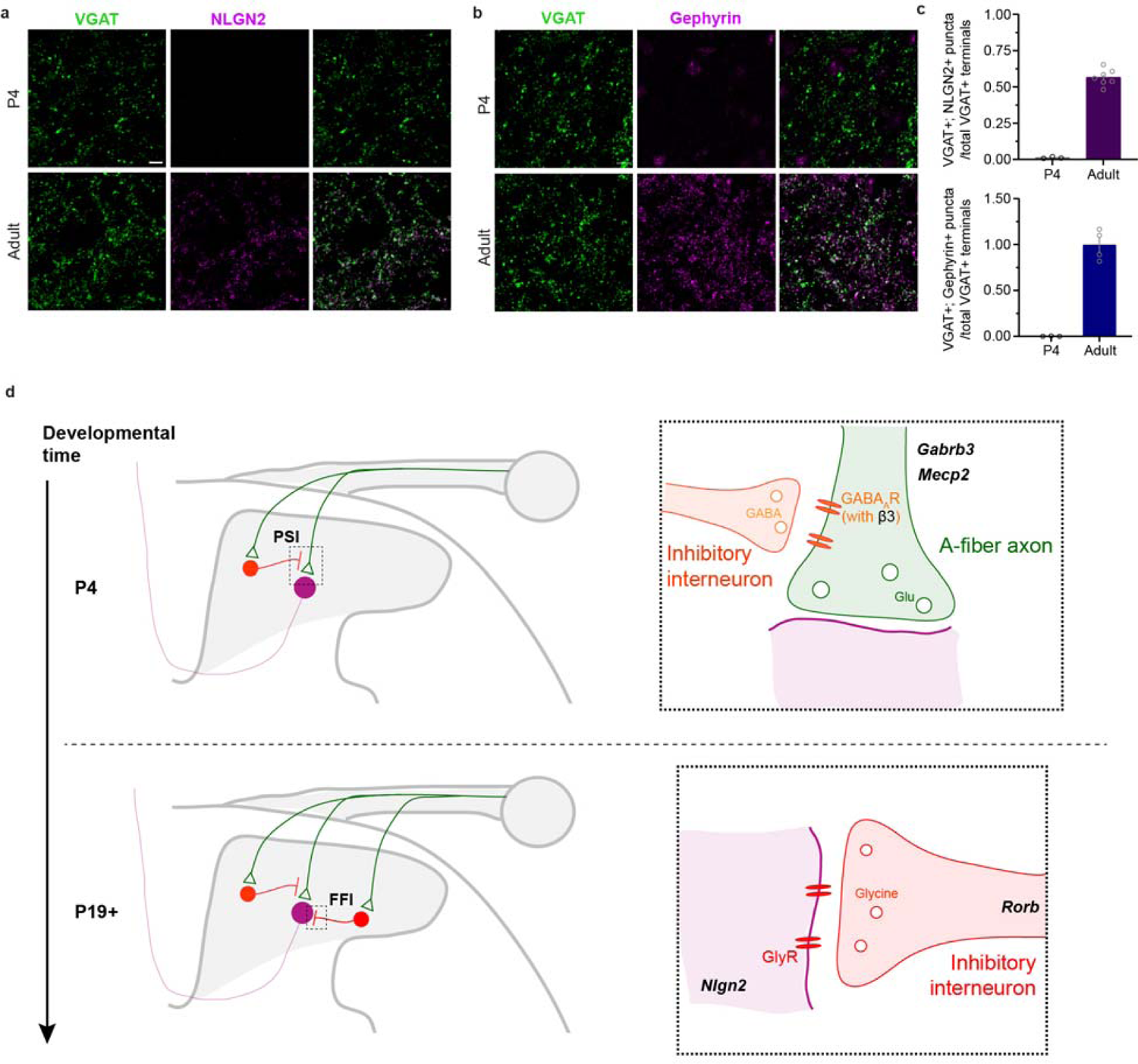
NLGN2 and gephyrin immunoreactivity are lacking at P4. Related to **Fig. 6**. **a**, **b,** Spinal cord immunohistochemistry of VGAT and NLGN2 (**a**) and gephyrin (**b**) in P4 and adult animals. Scale bar denotes 5 µm. **c**, Quantification of VGAT/NLGN2 (top) and VGAT/gephyrin (bottom) co-localization. **d**, Proposed model for the molecular and circuit development of presynaptic inhibition (PSI) and feedforward inhibition (FFI) in the spinal cord. Glu denotes glutamate, and GlyR denotes glycine receptor.

## METHODS

### Experimental model and subject details

All procedures performed in this study were approved by the Harvard Medical School Institutional Animal Care and Use Committee (IACUC). Male and female mice of mixed genetic backgrounds (C57BL/6J, 129/SvEv, CD1) were used for these studies. For adult behavioral testing, mice were weaned and ear notched for genotyping at P21 (+/- 2 days), and testing was done beginning at 6 weeks of age and complete by 8 weeks of age. For embryonic and neonatal behavior, mice were toe-clipped following testing and genotyped. For all behavioral assays, experiments were performed by investigators blinded to genotype. For electrophysiology experiments, mice were toe-clipped and genotyped prior to P4. Mice were group housed with littermates in standard housing on a 12-hour light/dark cycle.

### Mouse lines and genotyping

*Gabrb3* floxed mice were obtained from the Jackson Laboratory (008310) and were previously described^79^. *Gabrb3* Null mice were generated by crossing *Gabrb3* floxed mice to *E2a^Cre^* mice from the Jackson Laboratory (003724). The following primers were used to detect the wild-type, floxed, and null alleles: forward 5’-ATT CGC CTG AGA CCC GAC T-3’; reverse 5’-GTT CAT CCC CAC GCA GAC-3’; mutant reverse 5’-CCT TGG ACT GAG TCA CTG GAG-3’.

*Mecp2* Null mice were obtained from the Jackson Laboratory (003890) and were previously described^80^. The following primers were used to detect the wild-type and null alleles: common forward 5’-AAA TTG GGT TAC ACC GCT GA-3’; wild-type reverse 5’-CTG TAT CCT TGG GTC AAG CTG-3’; mutant reverse 5’-CCA CCT AGC CTG CCT GTA CT-3’.

*Nlgn2* floxed mice were obtained from the Jackson Laboratory (025544) and were previously described^33^. *Nlgn2* Null mice were generated by crossing *Nlgn2* floxed mice to *E2a^Cre^* mice and were subsequently backcrossed to C57Bl/6 mice for at least 6 generations. The following primers were used to detect the wild-type, floxed, and null alleles: forward 5’-CAA GCA CAG GGT TTC ACA GA-3’; mutant forward 5’-CACGAGGGCTTCAGTGTAGG-3’; reverse 5’-AGG CAA TGT GGT AGC TGG AG-3’.

*Rorb*^h1/h1^ mice were obtained from the Jackson Laboratory (006948) and were previously described^43^. The following primers were used to detect the wild-type and transgenic alleles: wild-type forward 5’-CTG TCC CTG TAT GCC TCT GG-3’; wild-type reverse 5’-AGA TGG AGA AAG GAC TAG GCT ACA-3’; transgene forward 5’-CTC AAC ACA CTA GAT GCC GAA G-3’; transgene reverse 5’-TGG CCA TCA CCA ACA ACA-3’.

*Nlgn4* Null mice were obtained from the MMRRC Repository (BayGenomics, 010566-UCD) and were previously described^36^. The following primers were used to detect the wild-type and null alleles: common forward 5’-CTT CCT ATC CTG TTA CTC TCA C-3’; wild-type reverse 5’-TAG GGA AAG CGG AAT TGA GTG TAA C-3’; mutant reverse 5’-ACA CTC CAA CCT CCG CAA ACT CCT-3’.

*Advillin^Cre^* mice were obtained from Fan Wang (Duke University) and were previously described^58^. The *Advillin^Cre^* transgene was identified using the following primers: 5′-CCC TGT TCA CTG TGA GTA GG −3′; reverse 5′-AGT ATC TGG TAG GTG CTT CCA G −3′; and internal control 5′-GCG ATC CCT GAA CAT GTC CAT C −3′.

*Cdx2^Cre^* mice were obtained from the Jackson Laboratory (009350) and were previously described^54^. The *Cdx2^Cre^* transgene was identified using the following primers: forward 5’-CTC GAC GTC TCC AAC CAT TG-3’; and reverse 5’-ATC TTC AGG TTC TGC GGG AA-3’.

*E2a^Cre^* mice were obtained from the Jackson Laboratory (003724) and were previously described^81^. *Emx1^Cre^* mice were obtained from the Jackson Laboratory (005628) and were previously described^82^. In both cases, the Cre transgene was identified using the following generic Cre primers: transgene forward 5’-CGG CAT GGT GCA AGT TGA AT-3’; transgene reverse 5’-AAC CAG CGT TTT CGT TCT GC-3’; internal positive control forward 5’-TTA CGT CCA TCG TGG ACA GC-3’; and internal positive control reverse 5’-TGG GCT GGG TGT TAG CCT TA-3’.

*Lbx1^Cre^* mice were previously described (MGI: 104867)^55^. The *Lbx1^Cre^* transgene was identified using the following primers: forward 5’-CGC CTT CCT CTC GCA CCG TC-3’; and reverse 5’-GGC AGC CCG GAC CGA C-3’.

*Ret^CreER^* animals were previously described (MGI: 97902)^50^. The *Ret^CreER^* transgene was identified using the following primers: common forward 5’-AGC GCA GGT CTC TCA TCA GT’; wild-type reverse 5’-GCA GGA GCA AAA TCA GCT TC-3’; and transgene reverse 5’-GCG CGC CTG AAG ATA TAG AA-3’.

*TrkB^CreER^* animals were previously described (MGI: 997384)^52^. The *TrkB^CreER^* transgene was identified using the following primers: TrkB CreER-1 (5’ UTR) 5’-GAC ACG CAC TCC GAC TGA-3’; TrkB CreER-2 (Intron 2): 5’-ACA CCT GCC TGA TTC CTG AG-3’; and TrkB CreER-3 (CreERT2) 5’-TCC TCA TCC TCT CCC ACA TC-3’.

*TrkC^CreER^* animals were previously described (MGI: 97385)^51^. The *TrkC^CreER^* transgene was identified using the following primers: forward 5’-TTT GCA GGT GTG TCC GTT TT-3’; and reverse 5’-GAC CGG TAA TGC AGG CAA AT-3’.

*Rosa26^LSL-synaptophysin-tdTomato^* (Ai34) mice were obtained from the Jackson Laboratory (012570). The *Rosa* locus was detected using the following primers: forward 5’-AAG GGA GCT GCA GTG GAG TA-3’; and reverse 5’-CCG AAA ATC TGT GGG AAG TC-‘3’. The presence of Ai34 was detected using: Ai forward 1 5’-GCT AAC CAT GTT CAT GCC TTC-3’; Ai forward 2 5’-GCT GAT CCG GAA CCC TTA AT-3’; and Ai34 reverse 5’-ATC TTG GTA GTG CCC CCT TT-3’.

*Rosa26^LSL-FSF-ReaChR-mCitrine^* mice were obtained from the Jackson Laboratory (024846). The *Rosa* site and CAG promoter were detected using the following primers: ROSA-CAG forward 5’-ACT TGC TCT CCC AAA GTC GCT CT-3’; ROSA-CAG reverse 5’-GCC AAG TGG GCA GTT TAC CG-3’; and ROSA-WT reverse 5’-TTT AAG CCT GCC CAG AAG ACT CCC-3’.

ReaChR expression was confirmed using ReaChR forward 5’-GAA GCG TTT CAT GAA TTT GAT AGC C-3’ and ReaChR reverse 5’-AGG CTG CTC TCG TAC TTA TCT TC-3’. CD1 animals were obtained from Charles River (#022) and C57Bl/J6 animals were obtained from the Jackson Laboratory (000664). *Advillin^FlpO^* mice were previously described^49^. Proper expression of each floxed allele using each Cre transgene was assessed using PCR. *Advillin^Cre^* and *Lbx1^Cre^* animals with post-Cre excision expression in tail biopsy tissue, or *Cdx2^Cre^*animals with excision in ear punch tissue were excluded from analyses.

### Tamoxifen treatment

Tamoxifen (Sigma T5648) was dissolved in 100% ethanol to 20 mg/mL, (30 minutes vortexing at room temperature) and then mixed with an equal volume of sunflower seed oil (Sigma S5007) and vacuum centrifuged for 30 minutes to evaporate the ethanol. When tamoxifen was delivered embryonically, progesterone (Sigma P0130) was also added to the ethanol at a concentration of 10 mg/mL. 20 mg/mL tamoxifen (with 10/ mg/mL progesterone) aliquots in sunflower seed oil were stored at −80LJC until the day of use, when they were thawed 15 minutes at room temperature, while protected from light. Pregnant mice received 3 mg tamoxifen by oral gavage at E11.5 (to label Aβ LTMRs using *Ret^CreER^*), E12.5 (to label Aβ SA-LTMRs using *TrkC^CreER^*), or E13.5 (to label Aβ RA- and Aδ-LTMRs using *TrkB^CreER^*).

### Behavioral testing

Male and female mice of mixed genetic backgrounds (C57BL/6J, 129/SvEv, CD1) were used for these studies. The only exceptions were *Nlgn2* and *Mecp2* germline mutant mice, which were backcrossed for at least 5 generations to a C57BL/6J background. For adult behavioral testing, littermates from the same genetic crosses were used as controls for each group (therefore, a combination of Cre-positive, floxed-negative and Cre-negative, floxed-positive mice were used for conditional knockout animal experiments). Prior to adult behavior testing, mice were weaned and ear notched (Kent Scientific, INS750075-5) for genotyping at P21 (+/- 2 days), and testing was done beginning at 6 weeks of age and complete by 8 weeks of age. For embryonic and neonatal behavior, mice were toe-clipped following testing and genotyped. For all behavioral assays, experiments were performed by investigators blinded to genotype. Mice were group housed with littermates in standard housing on a 12-hour light/dark cycle and free access to food and water. Cre alleles were always kept on the paternal genome, and all animals from each genetic cross had mothers of the same genotype, with the exception of crosses where the mother harbored either one or two floxed alleles (for example, *Gabrb3^flox/+^* or *Gabrb3^flox/flox^* mothers). No behavioral differences were observed between any wild-type, single or double flox, or Cre^+^ control groups. No differences between male and female mice of the same genotype were observed. Adult animals hemizygous for *Mecp2* were only compared to male control littermates. For adult behavioral testing, cages were changed once per week by an investigator and at least three days before the next behavioral assay. Prior to testing, animals were brought into the procedure room in their home cages and allowed to habituate for 30 minutes. All testing materials were cleaned thoroughly with 70% ethanol before and between trials.

### Open Field Test

Mice were placed into a 40 cm x 40 cm x 40 cm matte black acrylic testing chamber with matte white acrylic floor under dim lighting. Behavior was recorded from above for a duration of 10 minutes, and videos were tracked using custom MATLAB scripts to analyze distance traveled and time spent in center of arena. Time spent in center is measured as percent of time spent at a distance >5 cm from the edges of the chamber.

### Novel Object Recognition Test (NORT)

NORT testing was performed as previously described^23^. For two days, mice were individually habituated to the acrylic testing chamber used for the open field assay for 10 minutes each day under dim lighting. The next day, texture NORT was used as a measure of texture discrimination. During an initial learning phase, the mouse was placed into the testing chamber with two identical objects – either both “rough” or “smooth” plexiglass cubes (4 cm^3^), and visually identical – and allowed to freely explore the objects and chamber for 10 minutes. The mouse was then returned to its home cage for a 5 minute retention period, during which the chamber and objects were cleaned with 70% ethanol and one object was replaced with an object of novel texture (rough or smooth). The mouse was then placed back in the chamber for the 10 minute test phase. *Nlgn2* Null mice were subjected to a color/shape NORT, in which the objects were wooden blocks that differ in shape and color.

Video recordings were taken from above during both the learning and test phases. Custom MATLAB scripts tracked mouse position in the chamber, and time spent investigating objects in both phases was calculated. The preference for the novel object was calculated by measuring the time spent investigating the novel object divided by the time spent investigating both objects during the test phase. Mice that did not investigate both objects during the learning phase were excluded from analyses. Mice were whisker plucked three days before the start of habituation, which does not affect exploratory behaviors or total time spent investigating objects during NORT^23^.

### Prepulse Inhibition (PPI) of the Startle Reflex Assay

PPI was performed using the San Diego Instruments startle reflex system (SR-LAB Startle Response System) as described previously^23^. Mice were placed in a cylindrical enclosure just wide enough for the animal to turn around comfortably, within a soundproof chamber. A prepulse was delivered prior to an acoustic startle pulse (125 dB, 20 ms), and the mouse’s startle response was measured using an accelerometer mounted to the enclosure. For tactile PPI, the prepulse was a 0.9 PSI air puff (50 ms) delivered at variable interstimulus intervals (50, 100, 250, 500, and 1000 ms) prior to the acoustic startle pulse. Following an acclimation phase (5 minutes), testing sessions were arranged in blocks, consisting of 1) responses to acoustic pulse trials alone, 2) prepulse stimuli alone, and 3) pseudorandomized trials of prepulse/pulse (startle response), pulse alone (used for %PPI calculation), and no stimulation (to capture baseline movement). Inter-trial intervals varied from 10 to 30 seconds. Maximal accelerometer responses (in millivolts) were recorded in the 100 ms window following the end of each stimulus, and % PPI was calculated as 1-(startle response / pulse alone response). Response to air puff alone was measured as (prepulse alone/pulse alone response)-(no stimulation/pulse alone response). Animals whose baseline movement was greater than 25% of their acoustic startle response were not included in analyses (i.e. these animals exhibited insufficient acoustic startle amplitudes). All %PPI data reported in this study are from the 250 ms interstimulus interval trials.

### Elevated Plus Maze

The elevated plus maze consisted of four arms, each 30 cm long x 5 cm wide with white acrylic floors, and with two of the opposing arms having black acrylic walls that were 15 cm high. The maze stood on 40 cm tall legs. Testing was performed under dim lighting and video was recorded from above. The animal was placed at the center of the maze and allowed to explore for 10 minutes. The amount of time spent in each arm was tracked using a custom MATLAB script. The fraction time spent exploring the open arms was calculated as time spent in open arms / (time spent in open + closed arms).

### 3-Chamber Social Interaction Test

Sociability was assessed using the 3-chamber social interaction assay. Mice were first allowed to explore a three-chambered acrylic box with openings between the chambers for 5 minutes (each compartment was 20 cm wide x 40 cm long x 22 cm high). The outside walls of the chamber were black and opaque, and the two inner dividers were clear acrylic. After habituation, the test mouse was moved to the center chamber and clear partitions were put into place to block the entrances to the other two chambers. A novel mouse was then placed in a wire mesh cup in one chamber, while an identical, empty wire mesh cup (“object”) was placed on the other side. The partitions were then lifted, and the test mouse was free to explore for 10 minutes while video was recorded from above. The time spent in each of the three chambers during each test session was tracked using custom MATLAB scripts. The preference for novel mouse was calculated as the time spent investigating novel mouse / (time spent investigating novel mouse + object).

### Neonatal tactile sensitivity assay

All animals were group housed, with control and mutant animals in the same litters and cages. We did not observe any sex-related differences in P4 tactile reactivity in any genetic cross, so male and female mice of the same genotype were grouped together for final analyses. Male P4 animals hemizygous for *Mecp2* (*Mecp2*^-/y^) were compared to both male and female P4 control littermates.

For behavioral testing, P4 animals were removed from their home cage and brought into the behavioral testing room, and were returned to the home cage and litters immediately after testing. Animals were placed on warm, lightweight wool batting fabric, and the back of the mouse was positioned 2-3 mm below affixed air tubing. A camera (FLIR Integrated Imaging

Solutions, FL3-U3-13E4M-C) was mounted overhead and videos were acquired at 120 frames/second using FlyCapture SDK. Air puff stimuli of increasing intensity (0.10, 0.25, 0.50, 0.75, and 1.0 PSI, 50 ms each) were delivered sequentially with pseudo-randomized inter-stimulus intervals lasting between 20 and 30 seconds, using software and equipment from San Diego Instruments (SR-LAB Startle Response System). The 1.0 PSI stimulus was presented consecutively 10 times to test for average responsivity to the same stimulus and for habituation, totaling 14 trials (<7 minutes of testing) per animal. At the beginning of each session, an image of a ruler was used to ensure identical viewing distances across sessions. Videos were analyzed for peak body displacement 500 ms following the air puff stimulus using custom Python code. Trials in which the animal exhibited movement during the 10 frame baseline collection period prior to the stimulus were discarded during analysis. For habituation analyses, only animals that exhibited >20% of baseline movement to the 1^st^ presentation of the 1.0 stimulus were included for analysis. To calculate whether an animal exhibited habituation, the average displacement from the first three 1.0 PSI trials and average of the last three trials were compared, and animals exhibiting a >25% reduction in responsivity were considered to have habituated. The experimenter was blind to genotype of all test animals, as animals were toe clipped and genotyped after testing was completed.

For experiments related to Figure S2, 4% topical lidocaine (Henry Schein, 7280001) was applied to the back hairy skin of experimental animals following pre-treatment data collection and the animal was returned to its home cage for one hour before testing. For hair removal experiments, the back hairy skin was treated with commercial depilatory cream (NAIR, Church and Dwight Co.; Princeton, NJ) for 1 minute and then skin was gently cleaned with a dampened Kimwipe. The animal was then returned to its home cage for one hour before testing.

### Embryonic and neonatal optical reactivity assay

To measure optically-driven responses in ReaChR-labeled animals, a C-section was performed at embryonic day 18.5 and pups were allowed to recover on a warm pad for 15-30 minutes before behavioral testing. Animals were placed on a warmed, clear sheet of acrylic and a single 50 ms pulse of 470 nm light (1.1 mW/mm^2^; Thor Labs, M470F3) through a Ø400 µm Core, 0.39 NA patch cable (Thor Labs, M79L01) was used to deliver stimuli 5 times with 20-30 second ISI. The forepaw was tested first (light was directed to the glabrous side, from beneath), then the back hairy (at the nape of the neck, from above), then the hindpaw (same as forepaw). Either only the left or right paw was used for each animal. We report peak displacements in the 500 ms following stimulus onset, and average displacements were calculated as the average peak displacement to 5 optical stimuli. Following experiments, animals were cross-fostered to a CD1 female with an age-matched litter, and some animals were tested again at P0. Most P0 testing occurred in a separate set of animals that were delivered naturally or via C-section. No differences were observed between animals of the same genotype that were tested at both E18.5 and P0 versus only P0.

### Immunohistochemistry

For histology in wild-type animals, C57Bl6 animals were used for each experiment. For mutant versus control analyses, littermates were sacrificed in pairs or groups. Both males and females were used. For adult analyses, mice (P42-P84) were weighed, then anesthetized with isoflurane and transcardially perfused with 0.25 mL/g of body weight of Ames Media (Sigma, A1420) in 1xPBS with heparin (10 U/mL, Sigma, H3393-100KU), followed by 0.5 mL/g of 2% paraformaldehyde (PFA, Millipore Sigma, P6148) in PBS at room temperature (RT). No post-fixation was performed, and spinal cords and/or DRG were finely dissected from perfused mice.

For P0 and P4 spinal cord analyses in C57Bl6 animals, pups were deeply anesthetized on ice for several minutes before sacrificing. Mice were decapitated and the spinal column was dissected out, and drop-fixed in 2% PFA for 1 hour at RT. Tissue was then washed several times over 1 hour in 1xPBS. All spinal cord analyses were performed on lumbar segments (L3-L6).

Both adult and neonatal tissue was cryoprotected in 30% sucrose in 1XPBS for two nights at 4LJC, then embedded in NEG-50 (VWR, 84000-156) and frozen in tissue molds over a dry ice and ethanol slurry. Tissue was stored at −80LJC until transverse sections at 25 µm were taken on a cryostat, with littermate controls and mutants on the same slides, or C57Bl6 animals of different ages (P0, P4, adult) on the same slides. Slides were allowed to dry overnight at RT before staining, or stored at −20°C until staining.

After overnight drying or defrosting frozen samples for 15 minutes at RT, slides were rehydrated for 3×5 min in 1xPBS. For tdTomato and MeCP2 staining, slides were blocked for 2 h in 1x PBS containing 0.1% Triton X-100 and 5% normal goat or donkey serum (Jackson ImmunoResearch, 005-000-121 and 017-000-121). Sections were incubated with primary antibodies diluted in blocking solution (5% goat or donkey serum with no detergent) overnight at 4°C. Slides were washed 4×5 min in 1x PBS containing 0.02% Tween 20 (PBSt) and then incubated with species-specific secondary antibodies in blocking solution for 1 h at RT. Tissue was washed with PBSt 4×5 min and were mounted and coverslipped with Fluoromount Aqueous Mounting Medium with DAPI (Thermo Fisher, 00-4959-52). Slides were stored at 4°C.

For GABRB3, glycine receptor subunit α1, neuroligin-2, and gephyrin stains, a high salt protocol with antigen retrieval steps was used. High-salt PBS (HS-PBS) was comprised of 0.3M NaCl PBS. Tissue sections were incubated in 50% ethanol in ddH_2_O for 30 min to improve antibody penetration, then washed in HS-PBS for 3×10 min. Sections were then incubated at 37°C for 30 min in HS-PBS, then for 15 min in 0.001% trypsin + 0.001% CaCl_2_ at 37°C. Slides were washed in HS-PBS 3×10 min. Slides were incubated with primary antibodies diluted in HS-PBS with 0.3% Triton-X (HS-PBST) for 48 h at 4°C. After 48h, sections were incubated with primary antibodies for 30 min at RT. Sections were then washed in HS-PBS 6×10 min, and then incubated with secondary antibodies diluted in HS-PBST for 2 h at RT or overnight at 4°C. After secondary incubation, sections were washed in HS-PBST 3×10 min, and then mounted with Fluoromount Aqueous Mounting Medium with DAPI. Primary antibodies used: guinea pig anti-VGLUT1 (Millipore, AB5905, 1:1000), mouse anti-NLGN2 (Synaptic Systems, 129 511, 1:250), rabbit anti-GABRB3 (Biomatik, 1:500),^22^ rabbit anti-MeCP2 (Michael Greenberg, 1:1000), guinea pig anti-VGAT (Synaptic Systems, 131 004, 1:1000), mouse anti-glycine receptor α1 (Synaptic Systems, 146 111, 1:500), mouse anti-gephyrin (Synaptic Systems, 147 111, 1:500), and goat anti-mCherry (Sicgen, AB0040, 1:500). An array of goat or donkey derived Alexa Fluor 488, 546, and 647 conjugated secondary antibodies were used at a 1:500 dilution. In some experiments, the secondary antibody solution contained IB4 (Isolectin GS-IB4), Alexa 647 conjugate (Invitrogen, I32450) at 1:500 dilution.

For P0 glabrous forepaw and back hairy skin stains, pups were deeply anesthetized on ice for several minutes before sacrificing. Mice were decapitated and tissue samples were collected and drop-fixed in Zamboni fixative for 2 hours at 4°C. Tissue was then washed several times over 1 hour in 1xPBS. Tissue was cryoprotected in 30% sucrose in 1XPBS for two nights at 4LC, then embedded in Tissue-Tek O.C.T. Compound (Sakura, 4583) and frozen over a dry ice and ethanol slurry. Tissue was stored at −80°C until transverse sections at 30 µm were taken on a cryostat, and slides were allowed to dry overnight at RT before staining, or stored at −20LC until staining. Staining was performed using 5% normal goat serum protocol as described above. Primary antibodies used: rabbit anti-GFP (Abcam, ab6556, 1:500) and chicken anti-NFH (Aves Labs, NFH, 1:1000).

For immunohistochemistry following electrophysiological assays on 200 µm sagittal spinal cord slices, samples were incubated in 4% PFA for 30 minutes, washed several times in 1XPBS, and incubated overnight in IB4 in blocking solution (0.1% Triton-X and 5% NGS) at 4°C. Sections were then washed 3×5 minutes in 1XPBS and mounted on slides with Fluoromount Aqueous Mounting Medium with DAPI.

### Puncta Analysis

For puncta analyses, Z-stack images of spinal cord slices were taken on a Zeiss LSM 700 or LSM 900 confocal microscope using a 63X oil-immersion lens (Zeiss Plan-Apochromat 63X/NA 1.40). Images were taken in lamina III/IV of the dorsal horn, which in some experiments, was identified by immunostaining with IB4 to delineate lamina II_iv_. The depth of the z stack was determined by the expanse of VGLUT1+ or VGAT+ terminals. Imaging parameters were held constant across samples of each slide. Prior to colocalization analyses in C57Bl6 P0, P4, and adult animals, Contrast Limited Adaptive Histogram Equalization (CLAHE) was applied to each image and cross-correlation analyses were performed using custom MATLAB code to ensure comparability across experiments and conditions. Puncta analysis was performed in NIH ImageJ as described previously^22^. Briefly, a custom script created a mask of VGLUT1+ or VGAT+ terminals >0.5 µm in diameter, then performed thresholding on GABRB3, glycine receptor subunit α1, neuroligin-2, or gephyrin. Puncta of at least 0.1 µm in diameter that were contained within the VGLUT1 or VGAT mask were counted. Colocalization values are reported, for example for GABRB3, as GABRB3+, VGLUT1+ puncta/total number of VGLUT1+ per image. 3 images were taken per animal and an average colocalization value was taken per animal, which corresponds to each data point. The same parameters were used across tissue from the same slide.

### In situ hybridization

Detection of Nlgn2, Nefh, and Calca transcripts was performed by fluorescent in situ hybridization. Adult (>8 week old) animals were sacrificed by asphyxiation followed by cervical dislocation, and individual DRG ganglia were rapidly dissected from mice and frozen in dry-ice cooled 2 methylbutane and stored at −80LC until further processing. DRGs were cryosectioned at a thickness of 20 mm and RNA was detected using the RNAscope Fluorescent Multiplex Assay (ACD Bio) according to the manufacturer’s protocol. The following probes were used: Mm-Nlgn2 exons 3-5 (made-to-order, 300031), Mm-Nefh (443671), and Mm-Mrgprd (417921). Sections were mounted in Fluoromount Aqueous Mounting Medium (Thermo Fisher, 00-4958-02) and imaged on a Zeiss LSM 700 confocal microscope.

### Dorsal column and dorsal column nuclei injections

For PSDC labeling experiments for electrophysiology in mature animals, C57Bl6 male and female mice P14-P16 were anesthetized by isofluorane inhalation (1.5%–2.5%). Breathing rate and anesthesia were continuously monitored and isofluorane level adjusted as necessary. The back of the neck was shaved and then swabbed with betadine and 70% ethanol. A 5mm incision was made in the skin, and 0.5% lidocaine was applied to the incision site and underlying muscle. Muscles were separated or cut to expose the cervical vertebral column. The dura and arachnoid membranes between C1 and C2 were cut to expose the spinal cord. Six 50nL injections of Cholera Toxin Subunit B (Recombinant), Alexa Fluor™ 555 Conjugate (Fisher, C34776) mixed with a nominal volume of fast green were made bilaterally into the dorsal column under visual guidance with a glass pipette (three on each side), at a rate of 50 nL/minute using a Microinject system (World Precision Instruments). Injections were assessed by determining the extent to which the dorsal column took up the tracer, and an additional injection site was added when inefficient uptake was noted. Muscle and skin were then stitched together with sutures, and a liquid bandage (Nuskin) was applied. Prior to surgery, Nuskin was applied to the tail of the mother of experimental animals for habituation purposes. Buprenorphine was injected intraperitoneally for analgesia, and mice were returned to their home cage after recovery from anesthesia. The condition of the mice was monitored daily following surgery, until animals were sacrificed for electrophysiology experiments 3-5 days post-injection.

For PSDC labeling for P4 electrophysiological recordings, C57Bl6 P3 animals were initially anesthetized in an ice bath and hypothermia was maintained on a cold plate during the surgery. A 3mm incision was made in the skin, and 0.5% lidocaine was applied to the incision site and underlying muscle. Muscles were separated to expose the brainstem. The dura and arachnoid membranes were cut to expose the gracile and cuneate nuclei of the brainstem and 4-8 10 nL injections were made bilaterally at a rate of 100 nL/minute. Surgery was performed in under 10 minutes. Following surgery, mice were transferred to a sanitized, ventilated, and temperature-controlled 3×3×3 cm heated chamber (Warner, Dual Channel Temperature Controller, TC-344C) maintained at 34°C with nesting from home cage to minimize stress, and 4 mg/kg of carprofen was administered subcutaneously for analgesia. A cotton ball soaked with 3-5 mL of soymilk (Enfamil) was placed in the corner of the chamber, and animals were sacrificed no more than 12 hours following surgery for electrophysiological analyses.

### Spinal cord slice electrophysiological recordings

For electrophysiology in wild-type animals, C57Bl6 animals were used for each experiment. For mutant analyses, control littermates were used. P4 or P19-21 mice were deeply anesthetized (using ice or isofluorane, respectively) and rapidly transcardially perfused with RT choline chloride solution (92 mM choline chloride, 2.5 mM KCl, 1.2 mM NaH_2_PO_4_, 30 mM NaHCO_3_, 20 mM HEPES, 25 mM glucose, 5 mM sodium ascorbate, 2 mM thiourea, 3 mM sodium pyruvate, 10 mM MgSO_4_, 0.5 mM CaCl_2_). Vertebral columns were dissected out, and the lumbar spinal cord (L3-L6) was finely dissected out in ice cold choline chloride solution. Sagittal spinal cord sections (200 μm) were cut on a vibratome (Leica, VT1200S). Slices recovered for 30 minutes at 35°C in aCSF containing 2.0 mM CaCl_2_, 1 mM NaH_2_PO_4_, 119 mM NaCl, 2.5 mM KCl, 1.3 mM MgSO_4_, 26 mM NaHCO_3_, 25 mM dextrose, and 1.3 mM Na L-ascorbate (pH 7.4, 305-310 mOsm), oxygenated with 95% O2, 5% CO2 for 30 minutes. For mIPSC recording experiments, recovery aCSF included (*R*,*S*)-3-(2-Carboxypiperazin-4-yl)propyl-1-phosphonic acid (CPP, 5 μM; Abcam, ab120160) and 2,3-dihydroxy-6-nitro-7-sulfamoyl-benzo[f]quinoxaline (NBQX, 5 μM; Abcam, ab120046) to block excitatory glutamatergic activity in slices. For mEPSC recording experiments, recovery aCSF only included (R,S)-CPP to block NMDA receptors. After recovery period, slices were kept at RT in the recovery chamber until recording experiments began.

Slices were placed in the recording chamber and secured using a harp (Warner Instruments, 64-1421). Cells were visualized using infrared differential interference contrast (DIC) microscopy or fluorescent microscopy and recordings were performed in RT aCSF saturated with 95% O_2_/5% CO_2_ at a perfusion rate of 3-5 mL/min. Whole cell voltage-clamp recordings from lamina III and IV or from CTB-labeled spinal cord neurons were obtained under visual guidance using a 40x objective. Lamina III/IV was targeted based on distance from substantia gelatinosa, which is readily visualized under infrared DIC. Following break-in, cells were allowed to dialyze for 5 minutes prior to data collection.

For mIPSC recording experiments, aCSF contained 5 μM (R,S)-CPP, 5 μM NBQX, and 500 nM tetrodotoxin citrate (TTX; Tocris, 1069). For experiments in which GABAergic or glycinergic mIPSCs were isolated, 500 nM strychnine (Abcam, ab120416) or 5 μM SR 95531 (gabazine; Tocris, 1262), respectively, were either washed in or the entire recording was done with either drug in the aCSF. For mEPSC recording experiments, aCSF contained 5 μM (R,S)-CPP, 5 μM SR 95531, and 500 nM strychnine, and 500 nM TTX.

For mIPSC recordings, patch electrodes (3.0-4.5 MΩ) were filled with a CsCl-based internal solution containing 110 mM CsCl, 10 mM HEPES, 10 mM EGTA, 10 mM Cs-BAPTA, 4 mM CaCl_2_, 1 mM MgCl_2_, 10 mM TEA, 2 mM QX-314, and 0.2 mM D600 (pH 7.3, 295 mOsm) and neurons were voltage clamped at −60 mV. In some PSDC recording experiments, the internal solution also included ∼100 µM Alexa Fluor 488 for cell dialyzation and subsequent visualization. For mEPSC recordings, patch electrodes were filled with a CsGluconate-based internal solution containing 130 mM Cs-gluconate, 10 mM HEPES, 1 mM EGTA, 0.1 mM CaCl_2_, 5 mM TEA, 1 mM QX-314, and 0.2 D600 (pH 7.3, 295 mOsm) and neurons were voltage clamped at −60 mV. Data were acquired using a Multiclamp amplifier, a Digidata 1440A acquisition system, and pClamp10 software (Molecular Devices). Sampling rate was 20 kHz, and data were low-pass filtered at 3 kHz. No correction for junction potential was applied. Cells were discarded if residual uncompensated R_s_ was > 20 MΩ. Series resistance was monitored continuously throughout each experiment and cells were excluded from analysis if these values changed by more than 20% during the experiment. At least 100 events were analyzed per cell, which was performed in Clampfit (Molecular Devices).

### DRG electrophysiological recordings

P4 or P18-30 mice were deeply anesthetized and rapidly transcardially perfused with choline chloride solution as described above. The spinal column was removed and 4-6 lumbar level DRG were dissected and placed onto coverslips coated with Poly-L-Lysine (Sigma, P8920). DRG were allowed to recover for 8 minutes (P4) or 15 minutes (P18-30) in oxygenated aCSF solution at 35°C containing 0.01-0.02% Collagenase P (Sigma, 11213857001). RuBi-GABA (Fisher, 3400) stock solution was prepared by adding 10 μL DMSO before adding H_2_O as per manufacturer instruction to improve solubility. DRG were visualized using infrared DIC microscopy and recordings were performed in RT oxygenated aCSF as described above, with 100 μM of RuBi-GABA. Recordings were performed in the dark, but computer monitors were not covered with filters. Patch electrodes (7.5-10.5 MΩ) were filled with a KCl-based internal solution, containing 140 mM KCl, 10 mM HEPES, and 0.5 mM EGTA (pH 7.3, 295 mOsm) and recordings were performed in the current clamp configuration. Only medium- to large-diameter cells (>35 μm diameter, >55 pF capacitance)^59^ were analyzed with resting membrane voltage (RMP) of −60 to −70 mV, and cells were discarded if RMP became higher than −55 mV during the recording. Cells were dialyzed for 5 minutes after break-in, and excitability and action potential waveforms were first assessed using increasing steps of current (100 pA). To perform uncaging experiments, LED whole field illumination was used through a water immersion 40x objective, and maximum depolarization in response to a 5 ms uncaging pulse (473 nM, ∼5 mW) was recorded. Uncaging was performed 3 times per cell with 1 minute in between each trial. The peak depolarization in a 10 ms window following the light pulse was analyzed, and the average of all three traces is reported per animal. No differences were found between C57Bl/6 and *Gabrb3*^flox^, or Cre-negative animals with respect to DRG depolarization responses to GABA or intrinsic excitability.

### Quantification and statistical analyses

For all figures, data are expressed as the mean values ± standard error of the mean (SEM), unless noted otherise. For behavioral experiments, the number of animals per group used in each experiment are shown as individual datapoints in a graph. All behavioral and electrophysiology experiments were conducted using two or more cohorts of animals. Power analysis statistical tests were used to predetermine sample size for all behavioral and electrophysiology experiments. No statistical methods were used to predetermine sample sizes for immunohistochemical experiments.

All datasets were tested for normality using a Shapiro-Wilk test, and Bartlett’s test for equal variance was applied to all datasets. Comparisons between groups in all experiments were performed using unpaired, two-tailed or one-tailed Student’s t test (in the case of two groups distributed normally), Welch’s t test (in the case of two groups distributed normally, but with unequal variances), Mann-Whitney *U* test (for non-parametric data), one-way ANOVA (in the case of three or more groups distributed normally), two-way ANOVA (in the case of two groups with multiple conditions or time points), Kruskal-Wallis test (non-parametric test in the case of three or more groups), or Fisher’s Exact test (for comparing categorical variables). Comparisons between groups with significant differences are indicated above the appropriate groups. Asterisks indicate statistically significant *P* values for Student’s t tests, unless otherwise noted in the figure legends. For one-way ANOVAs, post hoc comparisons were performed using the post hoc test indicated in the figure legend. Statistically significant *P* values of post hoc comparisons are represented with asterisks in figures. ^#^*P*<0.1, \**P*<0.05; \*\**P*<0.001; \*\*\**P*<0.0001. All statistics were performed using GraphPad Prism.

### Data availability

All data generated during this study will be available by request.

### Code availability

Custom scripts used in this study will be available by request.

## ACKNOWLEDGEMENTS

We thank Michelle DeLisle for assistance with building the neonatal air puff system, Charalampia Koutsioumpa for help developing the optogenetic variant of the neonatal assay, Riana Pozsgai for assistance with mouse husbandry, Michael Greenberg for sharing the MeCP2 antibody, and Ofer Mazor and Pavel Gorelik of the Harvard Research Instrumentation Core for help with behavioral assay development. We thank members of the Ginty lab for helpful discussions and comments on the manuscript. This work was supported by NSF GRFPs DGE1745303 (AT and RH), a Mahoney Postdoctoral Fellowship (JT), a Gordon Postdoctoral Fellowship (JT), NIH R00NS101057 (LLO), NIH R01NS122788 (LLO), NIH R01NS126691 (LLO), the Hock E. Tan and K. Lisa Yang Center for Autism Research (DDG and LLO), NIH R35 5R35NS097344-05 (DDG), and the Edward R. and Anne G. Lefler Center for Neurodegenerative Disorders (DDG). LLO is a New York Stem Cell Foundation – Robertson Investigator; this research was supported by the New York Stem Cell Foundation. DDG is an investigator of the Howard Hughes Medical Institute. This article is subject to HHMI’s Open Access to Publications policy. HHMI lab heads have previously granted a nonexclusive CC BY 4.0 license to the public and a sublicensable license to HHMI in their research articles. Pursuant to those licenses, the author-accepted manuscript of this article can be made freely available under a CC BY 4.0 license immediately upon publication.

## SUPPORTING INFORMATION

**Supplementary Movie 1.** Displacement responses to 1.0 PSI air puff stimuli in control and mutant postnatal day 4 animals.

**Supplementary Movie 2.** Displacement responses to activation of low-threshold mechanoreceptors in control and mutant E18.5 animals.

## AUTHOR CONTRIBUTIONS

AT, LLO, and DDG conceived the study. AT and AA performed adult behavioral experiments and analyses. AT, IA, and AA performed immunohistochemistry experiments and analyses. AT and RH developed the embryonic/neonatal reactivity assays, RH created the analysis pipeline for these experiments, and AT performed the experiments. AT and IA did the dorsal column/dorsal column nuclei injections, with help from JT. JT developed the DRG electrophysiology preparation, and AT performed DRG and spinal cord electrophysiology experiments with help from JT. AT and DDG wrote the manuscript with input from all authors.

